# Dissecting the heterogeneous subcortical brain volume of Autism spectrum disorder (ASD) using community detection

**DOI:** 10.1101/2020.09.09.288993

**Authors:** Ting Li, Martine Hoogman, Nina Roth Mota, Jan K. Buitelaar, the ENIGMA-ASD Working Group, Alejandro Arias Vasquez, Barbara Franke, Daan van Rooij

**Author notes:** **Corresponding author**, Daan van Rooij, PhD, Department of Cognitive Neuroscience, Donders Institute for Brain, Cognition and Behaviour, Radboud University Medical Center, Kapittelweg 29, 6525 EN, Nijmegen, The Netherlands. Contributing members from the ENIGMA-ASD Working Group are (in alphabetical order): Adriana Di Martino, Alessandra Retico, Beatriz Luna, Bob Oranje, Celso Arango, Christine Deruelle, Christine Ecker, Christine M. Freitag, Clodagh M. Murphy, Damien Fair, Declan G.M. Murphy, Devon Shook, Eileen Daly, Evdokia Anagnostou, Fabio L.S. Duran, Fengfeng Zhou, Filippo Muratori, Geraldo F. Busatto, Gregory L. Wallace, Guillaume Auzias, Ilan Dinstein, Ilaria Gori, Jackie Fitzgerald, Jane McGrath, Jason Lerch, Jennifer Fedor, Joost Janssen, Joseph A. King, Katya Rubia, Kirsten O’Hearn, Liesbeth Hoekstra, Louise Gallagher, Luisa Lázaro, Mara Parellada, Margot Taylor, Maria Jalbrzikowski, Marlene Behrmann, Meiyu Duan, Michela Tosetti, Olga Puig, Pedro G.P. Rosa, Rosa Calvo, Sara Calderoni, Sarah Durston, Shlomi Haar, Stenfan Ehrlich.

## Abstract

Structural brain alterations found in Autism Spectrum Disorder (ASD) have previously been very heterogeneous, with overall limited effect sizes for every region implicated. In this study, we aimed at exploring the existence of subgroups in ASD, based on neuroanatomic profiles; we hypothesized that effect sizes of case/control difference would be increased in defined subgroups. Using the dataset from the ENIGMA-ASD Working Group (n=2661), exploratory factor analysis (EFA) was applied on seven subcortical volumes of individuals with ASD and controls to uncover the underlying organization of subcortical structures. Based on earlier findings in ADHD patients and controls as well as data availability, we focused on three age groups: boys (aged 4-14 years), male adolescents (aged 14-22 years), and adult men (aged >=22 years). The resulting factor scores were used in a community detection (CD) analysis, to cluster participants into subgroups. Three factors were found in each sample, with the factor structure in adult men differing from that in boys and male adolescents. From the patterns in these factors, CD uncovered four distinct communities in boys and three communities in adolescents and adult men, irrespective of ASD diagnostic status. The effect sizes of case/control comparisons appeared more pronounced than in the whole sample in some communities. Based on subcortical volumes, we succeeded in stratifying our participants into more homogeneous subgroups with similar brain structural patterns. The stratification enhanced our ability to observe case/control differences of subcortical brain volumes in ASD, and may help explain some of the heterogeneity of previous findings in ASD.

## Introduction

Autism spectrum disorder (ASD) is a neurodevelopmental disorder, which is characterized by persistent deficits in communication and social-emotional reciprocity combined with repetitive and stereotypical behaviors and interests [1]. The worldwide prevalence estimate for ASD is around 1,4% [2–4], with an estimated 3:1 higher prevalence rate in males than in females [5,6].

Structural brain alterations have been reported in ASD for several decades [7], with particularly lifespan-stable alterations observed especially in the subcortical areas, though existing literature shows considerable heterogeneity regarding the direction and size of subcortical alterations in ASD [8,9]. A number of studies have shown enlargement of amygdala, especially in children with ASD [10–14], while other studies on a wide age range of subjects reported either no differences [15–17] or a volumetric reduction of amygdala volume in ASD [18,19]. Findings from cross-sectional studies on hippocampal status have not reached consistency either. Increased and decreased hippocampal volumes have been found in ASD, irrespective of age [10,15,18,20,21]. Overall enlargement of the striatum in individuals with ASD has been reported compared to healthy controls [22–24]; however, notable inconsistencies also exist in this literature [9,25,26]. Similarly discrepant findings exist for the thalamus [24,25,27,28]. Recently, the ENIGMA-ASD Working Group conducted a large-scale case/control mega-analysis based on 51 existing datasets and reported individuals with ASD to have smaller subcortical volumes in the pallidum, putamen, amygdala, and nucleus accumbens [29]. However, all effect sizes observed within this large sample were small.

We expect that these limited effect sizes may be due to the heterogeneity of neuroanatomical profiles that exists within both the clinical and general population. Earlier clustering studies have shown that it was possible to stratify a population based on their neuroanatomical profiles, which increased the power to detect case/control differences within each subgroup [30]. Similarly, our recent findings from the ENIGMA-ADHD Working Group also showed distinct subgroups based on subcortical brain patterns present in male participants with and without ADHD [31]. Rather than expecting to find consistent anatomical alterations across the entire ASD population, it may therefore be more reasonable to first stratify both subjects with ASD and healthy controls into more homogeneous subgroups based on their neuroanatomical profiles, and subsequently investigate ASD diagnostic group differences within each subgroup.

Here, using subcortical brain volume data from the ENIGMA-ASD Working Group, we applied exploratory factor analysis (EFA) and community detection (CD) to explore the existence of more homogeneous subgroups in participants with and without ASD. We expected that similar subgroups should be observed within cases and controls, and that the effect sizes of case/control comparisons in these subgroups would be increased within each subgroup.

## Materials and methods

### Participants and ASD assessment

The analyzed magnetic resonance imaging (MRI) data in the current study come from the ENIGMA-ASD Working Group (http://enigma.ini.usc.edu/ongoing/enigma-asd-working-group). Full details about the international ENIGMA-ASD Working Group sample have been described before [29]. The working group implemented a data freeze in July 2018, at which point 1,353 patients with ASD and 1,308 healthy controls were included.

Based on earlier findings in subjects with ADHD, we expected sex difference in subcortical brain volumes organization [31], and given the limited data availability in females (only 145 girls, 45 female adolescents, and 33 women with ASD in ENIGMA-ASD cohort), we decided to only focus on male participants in the current study. We subdivided the full cohort into three subsamples based on age, a subsample comprised of 772 boys with ASD and 733 healthy controls (aged 4-13 years), a subsample of 360 male adolescents with ASD and 321 healthy controls (aged 14-22 years), and a subsample of 221 adult men with ASD and 254 healthy controls (aged >22 years). Information on the cohort and the subsamples in the current study is presented in **Table 1** and **Table S1**.

**Table 1:** Information on the three subsamples of the ENIGMA-ASD Working Group dataset.

| Variables | Boys |  | Male adolescents |  | Adult men |  |
| --- | --- | --- | --- | --- | --- | --- |
|  | Patients | Controls | Patients | Controls | Patients | Controls |
| **N** | 772 | 733 | 360 | 321 | 221 | 254 |
| **Mean Age (SD)** | 10.5 (2.8) | 10.6 (2.5) | 18.0 (2.0) | 17.9 (2.0) | 31.7 (9.4) | 30.7 (8.1) |
| **Mean IQ (SD)** | 103.9 (19.5) | 111.0 (15.5) | 105.4 (17.8) | 111.8 (12.4) | 109.7 (14.9) | 115.1 (11.6) |

*Note:* SD: Standard deviation; IQ: intelligence quotient

### Neuroimaging Segmentation

Structural T1-weighted brain MRI scans were collected at the various contributing sites. The MRI data were segmented using standardized ENIGMA imaging protocols based on FreeSurfer version 5.3 (http://enigma.ini.usc.edu/protocols/imaging-protocals/). Given the importance and stability of subcortical brain alterations in ASD as well as the need to limit the degrees of freedom to reach robust results, we selected the seven subcortical structures from the previous ENIGMA-ASD study for the current study. For each participant, the mean of the 7 subcortical volumes for two hemispheres were used for the analyses. The subcortical volumes were regressed with age, age^2, intracranial volume (ICV), and cohort site in the whole ENIGMA-ASD cohort for children and the rest of participants separately to allow for non-linear patterns of subcortical brain volumes across age.

### Factor Analysis

We performed exploratory factor analysis (EFA) to uncover the latent structure underlying the subcortical brain, and reduce the input variables to a more parsimonious model consisting of fewer factors than the total number of subcortical volumes. Following our previously established analysis pipeline [31], covariance matrices and squared multiple correlation were built as prior communality estimates for each subject over all subcortical volumes. Subsequently, maximum likelihood method and oblique rotation were applied to extract factors in the EFA. If the loading on the factor was 0.40 or more, a variable would be loaded on one factor. Model fitness was evaluated by Tucker Lewis Index (TLI), Bayesian information criterion (BIC), and the root mean square error of approximation (RMSEA). Given the EFA generated differential model outcome in the adult males as compared to the boys and adolescents, Confirmation Factor Analysis (CFA) was applied to test whether the factor structure generated in adult men was superior to that of the factor structure observed in the other two subsamples. This was done by evaluating Comparative Fit Index (CFI), TLI, BIC, and RMSEA between the resulting models. The analyses were conducted in R programming v3.6.2 using the ‘psych’ package.

### Community Detection (CD)

To identify distinct subgroups of participants based on factor scores generated from subcortical volumes, we utilized community detection (CD) [32,33]. Based on the normalized factor scores, n × n weighted, undirected networks were built to obtain distance information among participants. Then, we performed a weight-conserving modularity algorithm to identify distinct communities of participants in each network [30,33]. The algorithm sorts iteratively nodes (participants in this study) into communities until the modularity (Q) reaches maximum to find the optimal partition. The variation of information (VOI) was calculated to assess robustness of community structure. VOI indicates the variance between the original and perturbed networks over a range of alpha, which ranges between 0 and 1 [34]. The CD analyses were performed in Matlab [33].

### Statistical Analysis

Descriptive statistics of age and estimated intelligence quotient (IQ) were compared between individuals with and without ASD, using independent-samples t-test or Analysis of Covariance (ANCOVA). Chi-square test was used to check whether the distribution between communities differs for ASD cases and controls at each age bin. Within each sample, we compared subcortical factor scores and subcortical brain volumes between individuals with ASD and healthy controls using a t-test in each subgroup. Multivariate analysis of variance (MANOVAs) was applied to test which grouping (brain-based subgroup or ASD diagnosis group) showed a main effect on subcortical brain volumes in each subsample. False discovery rate (FDR) correction was used to correct for multiple comparisons of case-control differences within communities in the factor scores and subcortical volumes, separately in each age bin. All analyses were performed in IBM SPSS Statistics 25.

## Results

### 1. Participant characteristics

Demographic information about the three subsamples in current study is presented in **Table 1**. There were no case/control differences in age in each subsample after regressing the effect of cohort site (boys: t = −1.2, *P_adjusted_* = 0.46; male adolescents: t = 0.97, *p_adjusted_* = 0.53; adult men: t = 1.29, *p_adjusted_* = 0.42). Case/control differences in IQ were significant in each sample, with participants with ASD showing lower IQ than controls (Boys: F = 45.1, df = 1, *p_adjusted_* = 8.8e-10; male adolescents: F = 26.5, df = 1, *p_adjusted_* = 5.8e-06; adult men: F =17.7, df = 1, *p_adjusted_* = 2.6e-04).

### 2. EFA on subcortical volumes

#### 2.1 EFA in boys

EFA was applied to the residualized subcortical volumes in boys with and without ASD separately and together, which resulted in the similar factor structure. Three eigenvectors were extracted from the covariance matrix (Model fitness: TLI = 0.95, BIC = 1.94, RMSEA = 0.07). The first eigenvector is comprised of the volumes of caudate nucleus, globus pallidus, nucleus accumbens, and putamen. The second eigenvector included the hippocampus and amygdala, and the third eigenvector only included the thalamus. We interpreted them as ‘basal ganglia’, ‘limbic system’, and ‘thalamus’ (**Figure 1, Figure S1**). The three eigenvectors accounted for 30%, 16%, and 9% of the total shared variance, respectively.

**Figure 1:**
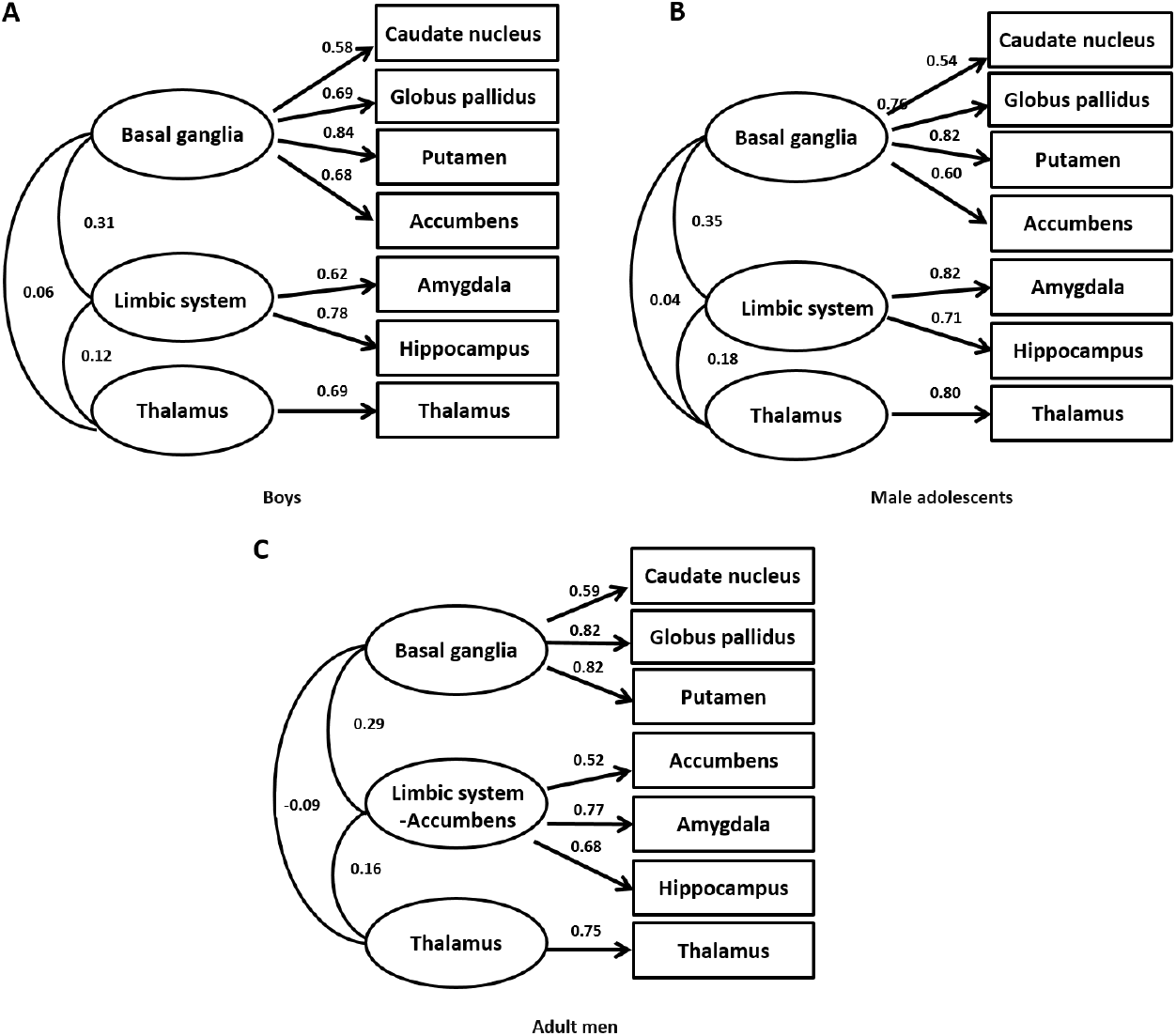
The three-factor model that was generated by EFA. A boys. B: male adolescents. C: adult men.

#### 2.2 EFA in male adolescents

EFA was next performed in male adolescents, including both participants with and without ASD. The same three eigenvectors as in boys were extracted (Model fitness: TLI = 0.94, BIC = −3.72, RMSEA = 0.08) (**Figure 1, Figure S1**). The proportion of variance accounted for by each eigenvector was 28%, 20%, and 12% of the total shared variance, respectively.

#### 2.3 EFA in adult men

In the subsample of adult men, EFA resulted in a different factor structure from that observed in boys and male adolescents (Model fitness: TLI = 1.01, BIC = −16.99, RMSEA = 0.00). The volumes of caudate nucleus, globus pallidus, and putamen loaded on the first eigenvector, which was interpreted as “basal ganglia”; The second eigenvector comprised the nucleus accumbens, hippocampus, and amygdala, which we named “limbic system-accumbens”; the third eigenvector only included the thalamus (**Figure 1, Figure S1**). The three eigenvectors accounted for 28%, 21%, and 12% of the total shared variance. The factor structure with nucleus accumbens loading on the second eigenvector was superior compared to the factor structure observed in boys and male adolescents (Model fitness: CFI= 0.70, TLI = 0.47, BIC = 48570.5, RMSEA = 0.24 compared to CFI= 0.59, TLI = 0.28, BIC = 48688.2, RMSEA = 0.28; chi square difference = 117.69, *p_adjusted_* = 1.1e-14).

### 3. CD in each sample based on subcortical factor scores

#### 3.1 CD in boys

The CD algorithm was first performed on the subcortical factor scores in boys (with and without ASD). Four distinct communities were generated, each comprising between 22.9% and 26.7% of the sample and containing boys with and without ASD (**Figure 2; Table 2**). Community 1 was characterized by increased volume of the basal ganglia and limbic system, but a smaller thalamus compared to the average volume of the whole sample. Community 2 showed a smaller basal ganglia and limbic system, but larger thalamus. Community 3 had a larger volume in the limbic system, but smaller basal ganglia, compared to the average volume. Community 4 had a larger basal ganglia, and smaller limbic system and thalamus compared to the average volume.

**Figure 2:**
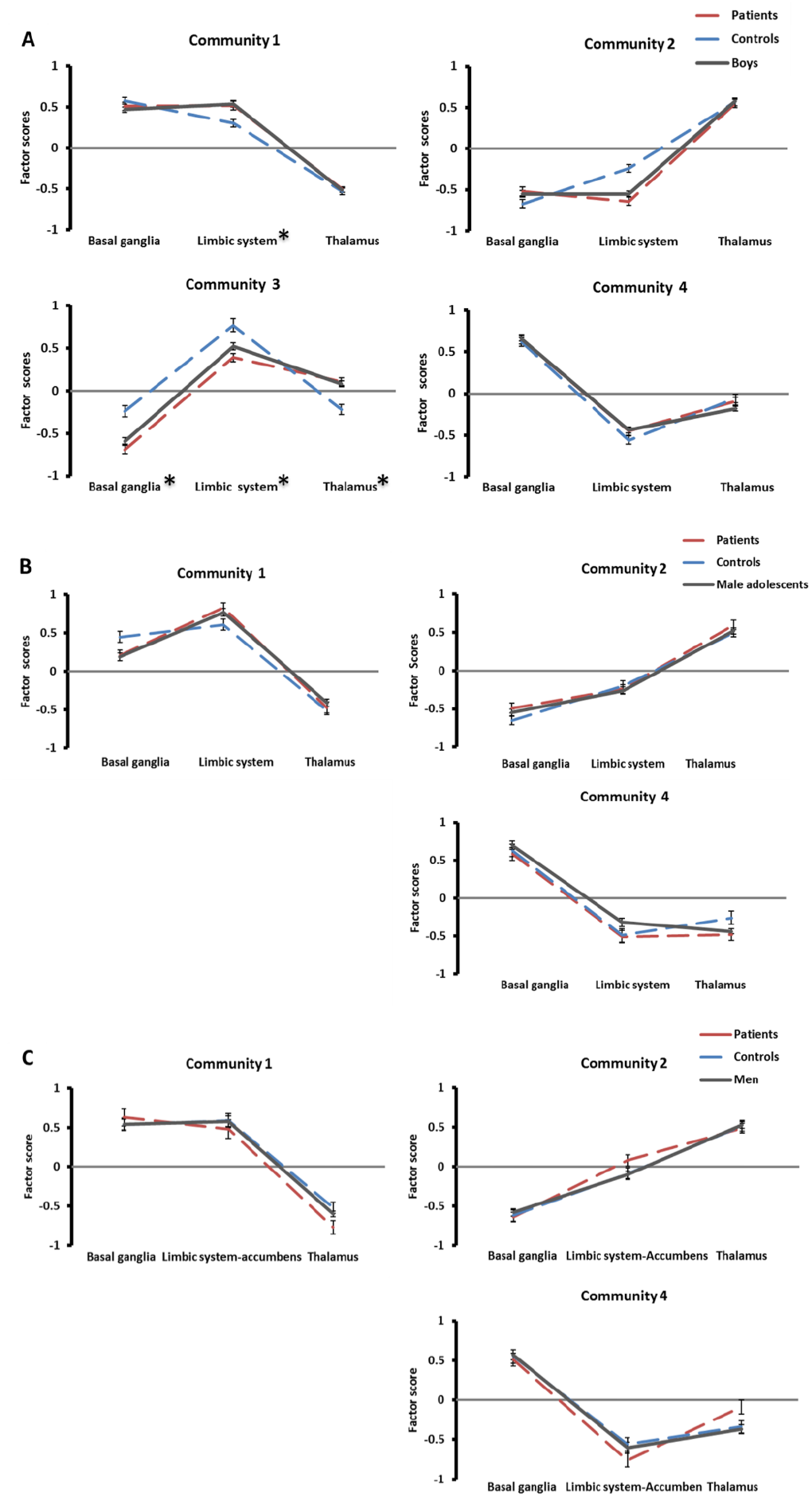
Subgroups generated by CD. A boys. B: male adolescents. C: adult men. ***Note:*** Lines represent participants in each community from CD. Y-axis indicates the mean factor scores for each factor. Error bars: standard error of the mean. *indicates case/control differences of subcortical factor scores are significant.

**Table 2:** The percentages of participants in each community of the three subsamples.

| **Sample** | **Total** | **Patients** | **Controls** |
| --- | --- | --- | --- |
| **Boys (N)** | **1505** | **772** | **733** |
| 1 | 381 (25.3%) | 221 (28.6%) | 200 (27.3%) |
| 2 | 402 (26.7%) | 204 (26.4%) | 240 (32.7%) |
| 3 | 345 (22.9%) | 193 (25.0%) | 129 (17.6%) |
| 4 | 377 (25.0%) | 154 (19.9%) | 164 (22.4%) |
| **Q values** | 0.45 | 0.46 | 0.43 |
| **Male Adolescent (N)** | **681** | **360** | **321** |
| 1 | 184 (27.0%) | 105 (29.2%) | 105 (32.7%) |
| 2 | 305 (44.8%) | 159 (44.2%) | 143 (44.5%) |
| 4 | 192 (28.2%) | 96 (26.7%) | 73 (22.7%) |
| **Q values** | 0.47 | 0.48 | 0.48 |
| **Men (N)** | **475** | **221** | **254** |
| 1 | 142 (29.9%) | 60 (27.1%) | 75 (29.5%) |
| 2 | 232 (48.8%) | 104 (47.1%) | 119 (46.9%) |
| 4 | 101 (21.3%) | 57 (25.8%) | 60 (23.6%) |
| **Q values** | 0.44 | 0.47 | 0.44 |

*Note:* Q values: the quality index of modularity

#### 3.2 CD in male adolescents

CD in male adolescents resulted in three communities. Each community accounted for 27.0% to 44.8% of the sample. No participants were present in the equivalent of Community 3 from the CD analysis in boys (**Figure 2, Table 2**). The three remaining communities had quite similar features to the equivalent communities in boys. Community 1 was characterized by increased volumes of the basal ganglia and limbic system above the average volume, but with a smaller thalamus. The volume of basal ganglia and limbic system were smaller than average, but the thalamus had a larger volume in Community 2. Community 4 showed a larger basal ganglia, but smaller limbic system and thalamus than average in the adolescents.

#### 3.3 CD in adult men

In adult men, CD revealed three communities with the proportion of participants from 21.3% to 48.8% of the sample. The equivalent of Community 3 in boys was absent (**Figure 2, Table 2**). In Community 1, the basal ganglia and limbic system-accumbens had increased volumes compared to the average level over all groups, but the thalamus was smaller. Community 2 had a reduced volume of the basal ganglia, but a larger thalamus than average. The volume of basal ganglia in Community 4 was increased compared to the average volume, but the limbic system-accumbens and thalamus were smaller than average.

In all three CD analyses, the quality index (**Table 2**) and VOIs (**Figure S2**) indicated that these communities significantly differed from random networks, and the networks were robust against chance variation. In this way, the VOI analysis can be viewed as an internal replication method, showing that the CD results do not change when a random part of the sample is perturbed. There were no significant differences in the distribution of ASD cases and controls between communities at each age bin (Boys: Chi square = 10.6, df = 3, *p_adjusted_* = 0.08; Male adolescents: Chi square = 3.3, df = 2, *p_adjusted_* = 0.41; Adult men: Chi square = 2.5, df = 2, *p_adjusted_* = 0.49). The distribution of cases and controls in each cohort is presented in **Table S2-4**.

### 4. Case/control comparison of subcortical factor scores in ASD

We examined whether individuals with ASD showed altered subcortical factor scores from healthy controls, first in each age group and then in each community separately. The results, as presented in **Table 2** and **Figure 2,** indicate that boys with ASD had smaller basal ganglia than healthy controls in Community 3 (t = −5.6, *p_adjusted_* = 1.0e-06, *d* = −0.63, 95% CIs [−0.86, −0.41]). For the limbic system, boys with ASD compared to healthy controls showed increased volume in Community 1 (t = 3.1, *p_adjusted_* = 0.01, *d* = 0.30, 95% CIs [0.11, 0.49]), but reduced volumes in Community 2 and 3 (Community 2: t = −5.9, *p_adjusted_* = 1.4e-07, *d* = - 0.56, 95% CIs [−0.75, −0.37]; Community 3: t = −4.4, *p_adjusted_* = 1.6e-04, *d* = −0.50, 95% CIs [−0.73, −0.27]). In Community 3, boys with ASD had a larger thalamus than healthy controls (t = 4.5, *p_adjusted_* = 1.3e-04, *d* = 0.51, 95% CIs [0.28, 0.74]). In the sample of male adolescents and adult men, two case/control differences were found each, but did not survive FDR correction.

In **Table S5-S7**, we present case/control comparisons for each individual subcortical brain volume in the whole sample and each community. We observed several significant case/control differences within communities: eight case/control comparisons in boys and three in male adolescents survived FDR correction. The effect sizes ranged from *d* = −0.84 (95% CIs [−1,07, −0.60]) to *d* = 0.37 (95% CIs [0.14, 0.59]) within communities, which were more pronounced than those in the whole subsample in which effect sizes *d* ranged −0.29 (95% CIs [−0.44, −0.13]) to 0.04 (95% CIs [-0.14, 0.22]). MANOVAs indicated that the communities accounted for more variance in subcortical brain volumes than ASD diagnosis in each subsamples (Boys: Communities: F(21,4467) = 147.8, *padjusted* = 1.1e-14; ASD diagnosis: F(7,1487) = 0.95, *padjusted* = 0.69; Male adolescents: Communities: F(14, 1332) = 113.7, *padjusted* = 1.1e-14; ASD diagnosis: F(7, 665) = 4.38, *p_adjusted_* = 6.5e-04; Men: Communities: F(14, 920) = 3.12, *p_adjusted_* = 6.4e-04; ASD diagnosis: F(7,459) = 0.83, *p_adjusted_* = 0.74).

## Discussion

In this study, we aimed to dissociate subgroups of ASD participants based on neuroanatomic profiles of subcortical structures. We hypothesized that effect sizes of case/control differences would be larger within each subgroup. In our exploratory factor analysis (EFA), we found that the latent structure of subcortical volumes is comprised of three factors, which remain largely stable across the lifespan and are identical in those with and without ASD. Among them, we discerned four distinct communities in boys and three in male adolescents and adult men. Within several of the communities, effect sizes of case/control differences in neuroanatomical volume were much stronger than the average differences across the whole sample.

In the samples of boys and male adolescents, the same three-factor structures - basal ganglia, limbic system, and thalamus were observed based on their subcortical brain volume distribution in healthy controls and participants with ASD taken together. In adult men, the three-factor structure was slightly different; nucleus accumbens loaded onto the second factor, which we named ‘limbic system-accumbens’, instead of the limbic system factor. These structural patterns of subcortical brain volumes were found regardless of diagnostic status in those with and without ASD, which indicates that no qualitative differences in subcortical brain organization exist in ASD. The factor structures are largely in line with previous smaller scale studies looking at subcortical brain organization. One previous study using 322 healthy adults (age range 65-85 years) reported three clusters based on cortex and subcortical structures, with one cluster comprising of basal ganglia (caudate, putamen, and pallidum) and a second cluster including nucleus accumbens, amygdala, hippocampus and thalamus; cortical lobes were in the third cluster [35]. A study on 404 healthy adults (age range 51-59 years) indicated that subcortical brain volumes could be partitioned into three factors: basal ganglia/thalamus, nucleus accumbens, and a limbic factor [36]. In a recent study of the ENIGMA-ADHD Working Group, we found identical subcortical factor structure as in the current analysis - basal ganglia, limbic system and thalamus - existed in boys and adult men, which was irrespective of ADHD diagnosis and age [31]. Nucleus accumbens receives direct glutamatergic inputs from amygdala and hippocampus, and the nucleus accumbens shell may be regarded as a part of the extended amygdala [37]; this may explain why the nucleus accumbens loads on either the basal ganglia or the limbic factor in the current study. The variation of the factor structure between age groups observed in the current study, in which we used a lifespan approach, may suggest that the correlation between subcortical structures changes slightly during maturation, as has also been suggested previously [38].

Using CD analysis, each of the three subsamples could be stratified into similar subgroups with more homogeneous neuroanatomic patterns. Four communities were observed in boys, three were seen in the samples of male adolescents and adult men, irrespective of ASD status and age; The CD results indicated that the heterogeneity in subcortical brain volumes is nested within normative variability, with different neuroanatomic communities existing in both controls and patients [39]. Importantly, the observed community structure is highly consistent with our recent findings in the ENIGMA-ADHD Working Group [31]. The fact that we observe not only a similar factor structure, but also similar community structure in that sample greatly supports the robustness of our current analysis. In fact, the CD results in the ENIGMA-ADHD control group can be viewed as an independent, external validation of the currently observed community structure. This also allows us to investigate where subjects with ADHD and ASD show differences in their community structure. In the current analysis, Community 3 disappeared in adolescents and men. In the ENIGMA-ADHD analysis, we also observed that Community 3 was absent in healthy men, but not in men with ADHD. This reduction of subgroups from four in the subsample of boys to three in the male adolescents and adult men may be related to structural brain maturation over age, leading to less diversity in the organization of subcortical volumes in the population [40].

In the current study, analyzing case/control differences within communities indicated substantially larger effect sizes as compared to the previous study on the entire sample without stratification [29]; interestingly, case/control differences are not consistently present in each factor in each community. For example, boys with ASD have increased volume of the limbic system in Community 1, but smaller volume in Community 2 and 3 compared to healthy controls. The substantially larger effect sizes within subgroups suggest that neuroanatomically based subgroups may exist within the entire population, and that distinct/alternative ASD-related anatomical alterations may be present in different subgroups. An important consequence of these findings is that there might not be a single neuroanatomical risk profile for ASD. Instead, the altered brain structures associated with ASD may be dependent on both the age and the neuroanatomical subgroup of an individual. The results also may explain some subcortical heterogeneity found in previous studies, as previous smaller studies may have accidentally recruited a disproportionately higher number of any specific subgroup, which may result in observed contradictory subcortical alterations in ASD [41]. In the current study, the brain-based ASD subgroups accounted for more variance of subcortical brain volumes than just the ASD diagnostic groups. However, because we did not have available to us deep phenotypic information, we could not further characterize the clinical presentation of our brain-based subgroups. Therefore, we cannot entirely rule out the existence of confounding factors that may be related to the neuroanatomical profiles observed in the different communities.

This work has to be viewed in light of several strengths and limitations. Using the MRI dataset from the ENIGMA-ASD Working Group, we had a large sample size, which allowed us to explore underlying structural pattern and subgroups in ASD across the lifespan; however, as mentioned previously, the limited availability of demographic information precluded our ability to explore whether brain-based communities are linked to the clinical presentation of ASD. Moreover, in the current study, we only had sufficient power to run the analysis in male participants. Previous studies have reported sex differences in subcortical brain volumes [42], and different underlying subcortical organizations were reported in females from ENIGMA-ADHD cohort [31]. Given that sex-based differences in neuroanatomy are a central topic in ASD [43,44], further analysis including females may help us elucidate the association between neuroanatomical organization and the specific etiology of ASD in females.

In conclusion, using subcortical brain volume data from the ENIGMA-ASD Working Group, we were able to stratify subjects with and without ASD into more homogeneous subgroups based on underlying neuroanatomic organization. Our results indicate that this stratification may enhance our ability to observe case/control differences and may explain some of the contradictory results observed in previous, smaller studies of brain structure in ASD.

**Figure 3:**
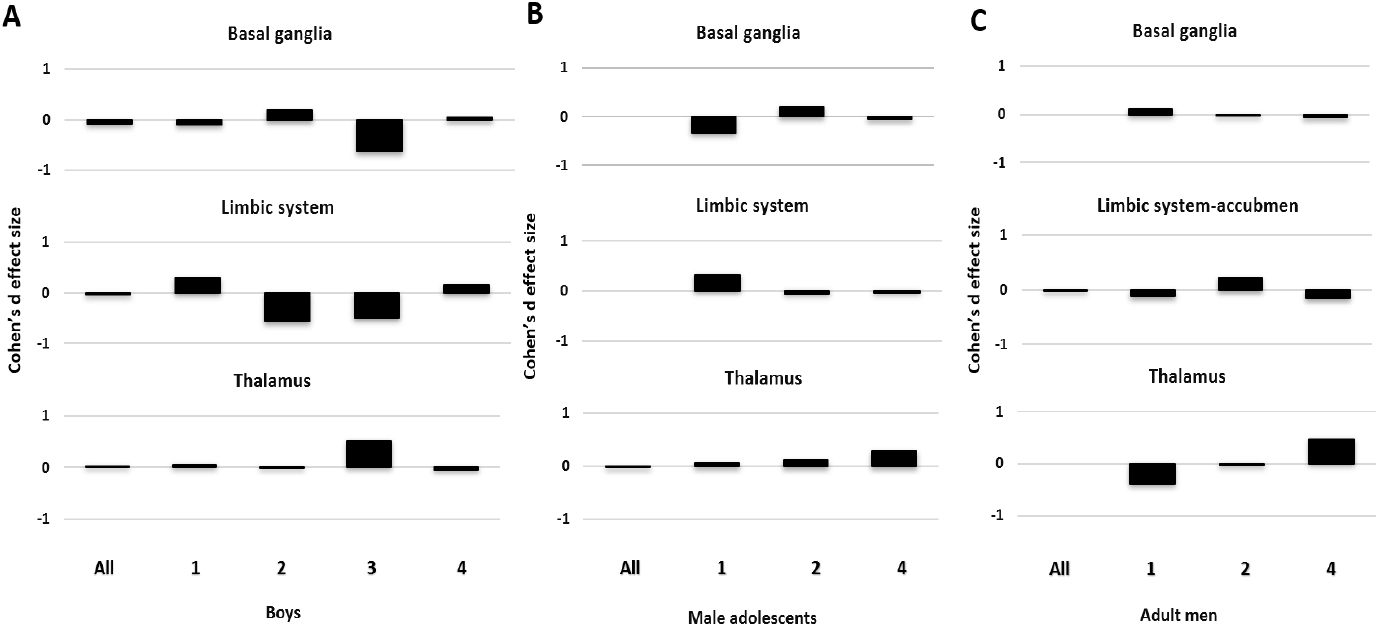
Effect sizes of case/control comparison within each community and the whole subsample. A boys. B: male adolescents. C: adult men. ***Note***: All: the whole subsample, 1: Community 1; 2: Community 2; 3: Community 3; 4: Community 4.

**Table 3:** Case/control Comparison of subcortical factor scores in ASD.

| Community | Basal ganglia |  |  |  |  | Limbic system |  |  |  |  | Thalamus |  |  |  |  |
| --- | --- | --- | --- | --- | --- | --- | --- | --- | --- | --- | --- | --- | --- | --- | --- |
|  | Mean factor scores |  | *p* value | *adjusted p* value | Cohen's d (95% CIs) | Mean factor scores |  | *p* value | *adjusted p* value | Cohen's d (95% CIs) | Mean factor scores |  | *p* value | *adjusted p* value | Cohen's d (95% CIs) |
|  | Patients | Controls |  |  |  | Patients | Controls |  |  |  | Patients | Controls |  |  |  |
| **Boys** | -0.03<br>(0.93) | 0.03<br>(0.91) | 0.15 | 0.37 | -0.07<br>(-0.18 - 0.03) | -0.01<br>(0.87) | 0.01<br>(0.85) | 0.54 | 0.74 | -0.03<br>(-0.13 - 0.07) | 0.01<br>(0.75) | -0.01<br>(0.78) | 0.57 | 0.74 | 0.03<br>(-0.07 - 0.13) |
| **1** | 0.51<br>(0.73) | 0.58<br>(0.58) | 0.33 | 0.53 | -0.10<br>(-0.29 - 0.10) | 0.52<br>(0.77) | 0.30<br>(0.67) | **2.0e-03** | **0.01** | 0.30<br>(0.11 - 0.49) | -0.50<br>(0.64) | -0.53<br>(0.58) | 0.41 | 0.62 | 0.05<br>(-0.14 - 0.24) |
| **2** | -0.52<br>(0.75) | -0.67<br>(0.78) | **0.04** | 0.15 | 0.19<br>(-0.01 - 0.38) | -0.64<br>(0.70) | -0.24<br>(0.71) | **6.6e-09** | **1.4e-07** | -0.56<br>(-0.75 - -0.37) | 0.55<br>(0.70) | 0.57<br>(0.70) | 0.82 | 1.0 | -0.02<br>(-0.21 - 0.17) |
| **3** | -0.69<br>(0.68) | -0.24<br>(0.75) | **5.2e-08** | **1.0e-06** | -0.63<br>(-0.86 - -0.41) | 0.39<br>(0.69) | 0.77<br>(0.86) | **1.5e-05** | **1.7e-04** | -0.50<br>(-0.73 - -0.27) | 0.11<br>(0.62) | -0.22<br>(0.68) | **1.1e-05** | **1.3e-04** | 0.51<br>(0.28 - 0.74) |
| **4** | 0.65<br>(0.69) | 0.62<br>(0.68) | 0.67 | 0.81 | 0.05<br>(-0.17 - 0.27) | -0.45<br>(0.60) | -0.56<br>(0.64) | 0.26 | 0.31 | 0.17<br>(-0.05 - 0.40) | -0.09<br>(0.56) | -0.06<br>(0.65) | 0.64 | 0.81 | -0.05<br>(-0.27 - 0.17) |
| **Male adolescents\*** | 0.00<br>(0.91) | 0.00<br>(0.93) | 1.0 | 1.0 | 5.7e-18<br>(-0.15 - 0.15) | 0.00<br>(0.91) | 0.00<br>(0.90) | 1.0 | 1.0 | 2.2e-16<br>(-0.15 - 0.15) | 0.00<br>(0.95) | 0.00<br>(0.82) | 1.0 | 1.0 | -6.5e-17<br>(-0.15 - 0.15) |
| **1** | 0.21<br>(0.68) | 0.45<br>(0.75) | **0.02** | 0.09 | -0.33<br>(-0.60 - -0.05) | 0.83<br>(0.66) | 0.61<br>(0.72) | **0.02** | 0.10 | 0.32<br>(0.04 - 0.59) | -0.47<br>(0.77) | -0.51<br>(0.57) | 0.68 | 0.81 | 0.06<br>(-0.21 - 0.33) |
| **2** | -0.49<br>(0.82) | -0.65<br>(0.69) | 0.07 | 0.22 | 0.21<br>(-0.02 - 0.44) | -0.24<br>(0.73) | -0.20<br>(0.85) | 0.06 | 0.20 | -0.06<br>(-0.28 - 0.17) | 0.60<br>(0.82) | 0.50<br>(0.72) | 0.28 | 0.48 | 0.12<br>(-0.10 - 0.35) |
| **4** | 0.58<br>(0.82) | 0.63<br>(0.76) | 0.67 | 0.81 | -0.06<br>(-0.37 - 0.24) | -0.51<br>(0.79) | -0.49<br>(0.75) | 0.89 | 0.94 | -0.03<br>(-0.33 - 0.28) | -0.48<br>(0.74) | -0.26<br>(0.75) | 0.06 | 0.20 | 0.29<br>(-0.60 - 0.01) |

| Community | Basal ganglia |  |  |  |  | Limbic system-Accumbens |  |  |  |  | Thalamus |  |  |  |  |
| --- | --- | --- | --- | --- | --- | --- | --- | --- | --- | --- | --- | --- | --- | --- | --- |
|  | Mean factor scores |  | *p*<br>*value* | *adjusted*<br>*p value* | Cohen's d<br>(95% CIs) | Mean factor scores |  | *p*<br>*value* | *adjusted*<br>*p value* | Cohen's d<br>(95% CIs) | Mean factor scores |  | *p*<br>*value* | *adjusted*<br>*p value* | Cohen's d<br>(95% CIs) |
|  | Patients | Controls |  |  |  | Patients | Controls |  |  |  | Patients | Controls |  |  |  |
| **Adult Men \*** | 0.00<br>(0.92) | 0.00<br>(0.93) | 1.0 | 1.0 | 2.0e-17<br>(-0.18 - 0.18) | 0.00<br>(0.89) | 0.00<br>(0.89) | 1.0 | 1.0 | -2.7e-16<br>(-0.18 - 0.18) | 0.00<br>(0.83) | 0.00<br>(0.82) | 1.0 | 1.0 | 6.9e-17<br>(-0.18 - 0.18) |
| **1** | 0.63<br>(0.88) | 0.54<br>(0.71) | 0.52 | 0.72 | 0.11<br>(-0.23 - 0.46) | 0.48<br>(0.95) | 0.59<br>(0.80) | 0.49 | 0.69 | -0.12<br>(-0.46 - 0.22) | -0.77<br>(0.66) | 0.52<br>(0.60) | **0.02** | 0.10 | -0.40<br>(-0.75 - -0.05) |
| **2** | -0.64<br>(0.60) | -0.62<br>(0.76) | 0.81 | 0.90 | -0.03<br>(-0.30 - 0.23) | 0.08<br>(0.75) | -0.09<br>(0.84) | 0.11 | 0.30 | 0.22<br>(-0.05 - 0.48) | 0.49<br>(0.63) | 0.52<br>(0.52) | 0.08 | 0.24 | -0.04<br>(-0.31 - 0.22) |
| **4** | 0.51<br>(0.62) | 0.55<br>(0.65) | 0.70 | 0.83 | -0.06<br>(-0.42 - 0.29) | -0.66<br>(0.66) | -0.56<br>(0.64) | 0.40 | 0.61 | -0.15<br>(-0.52 - 0.21) | -0.09<br>(0.68) | -0.39<br>(0.63) | **0.02** | 0.08 | 0.46<br>(0.09 - 0.82) |

**Note:** Adjusted p value : adjusted p value: FDR-correction in factor scores across age groups. Significant difference in bold. 95% CIs: 95% Confidence intervals. \* Community 3 is absent in male adolescents and adult men, because no healthy controls were loaded using CD.

## References

1 APA. American Psychiatric Association. Diagnostic and Statistical Manual of Mental Disorders, 5th edition. Arlington, VA.; 2013.

2 Baird G, Simonoff E, Pickles A, Chandler S, Loucas T, Meldrum D, et al. Prevalence of disorders of the autism spectrum in a population cohort of children in South Thames: the Special Needs and Autism Project (SNAP). Lancet. 2006;368(9531):210–5.

3 Christensen DL, Baio J, Van Naarden Braun K, Bilder D, Charles J, Constantino JN, et al. Prevalence and Characteristics of Autism Spectrum Disorder Among Children Aged 8 Years--Autism and Developmental Disabilities Monitoring Network, 11 Sites, United States, 2012. MMWR Surveill Summ. 2016;65(3):1–23.

4 Elsabbagh M, Divan G, Koh Y-J, Kim YS, Kauchali S, Marcín C, et al. Global prevalence of autism and other pervasive developmental disorders. Autism Res. 2012;5(3):160–79.

5 Kim YS, Leventhal BL, Koh YJ, Fombonne E, Laska E, Lim EC, et al. Prevalence of autism spectrum disorders in a total population sample. The American journal of psychiatry. 2011;168(9):904–12.

6 Loomes R, Hull L, Mandy WPL. What Is the Male-to-Female Ratio in Autism Spectrum Disorder? A Systematic Review and Meta-Analysis. Journal of the American Academy of Child and Adolescent Psychiatry. 2017;56(6):466–74.

7 Amaral DG, Schumann CM, Nordahl CW. Neuroanatomy of autism. Trends in neurosciences. 2008;31(3):137–45.

8 Donovan APA, Basson MA. The neuroanatomy of autism - a developmental perspective. J Anat. 2017;230(1):4–15.

9 Haar S, Berman S, Behrmann M, Dinstein I. Anatomical Abnormalities in Autism? Cerebral Cortex. 2014;26(4):1440–52.

10 Groen W, Teluij M, Buitelaar J, Tendolkar I. Amygdala and hippocampus enlargement during adolescence in autism. Journal of the American Academy of Child and Adolescent Psychiatry. 2010;49(6):552–60.

11 Munson J, Dawson G, Abbott R, Faja S, Webb SJ, Friedman SD, et al. Amygdalar volume and behavioral development in autism. Archives of general psychiatry. 2006;63(6):686–93.

12 Nordahl CW, Scholz R, Yang X, Buonocore MH, Simon T, Rogers S, et al. Increased rate of amygdala growth in children aged 2 to 4 years with autism spectrum disorders: a longitudinal study. Archives of general psychiatry. 2012;69(1):53–61.

13 Schumann CM, Hamstra J, Goodlin-Jones BL, Lotspeich LJ, Kwon H, Buonocore MH, et al. The amygdala is enlarged in children but not adolescents with autism; the hippocampus is enlarged at all ages. The Journal of neuroscience: the official journal of the Society for Neuroscience. 2004;24(28):6392–401.

14 Sparks BF, Friedman SD, Shaw DW, Aylward EH, Echelard D, Artru AA, et al. Brain structural abnormalities in young children with autism spectrum disorder. Neurology. 2002;59(2):184–92.

15 Barnea-Goraly N, Frazier TW, Piacenza L, Minshew NJ, Keshavan MS, Reiss AL, et al. A preliminary longitudinal volumetric MRI study of amygdala and hippocampal volumes in autism. Progress in neuro-psychopharmacology & biological psychiatry. 2014;48:124–8.

16 Haznedar MM, Buchsbaum MS, Wei TC, Hof PR, Cartwright C, Bienstock CA, et al. Limbic circuitry in patients with autism spectrum disorders studied with positron emission tomography and magnetic resonance imaging. The American journal of psychiatry. 2000;157(12):1994–2001.

17 Schumann CM, Amaral DG. Stereological analysis of amygdala neuron number in autism. The Journal of neuroscience: the official journal of the Society for Neuroscience. 2006;26(29):7674–9.

18 Aylward EH, Minshew NJ, Goldstein G, Honeycutt NA, Augustine AM, Yates KO, et al. MRI volumes of amygdala and hippocampus in non-mentally retarded autistic adolescents and adults. Neurology. 1999;53(9):2145–50.

19 Nacewicz BM, Dalton KM, Johnstone T, Long MT, McAuliff EM, Oakes TR, et al. Amygdala volume and nonverbal social impairment in adolescent and adult males with autism. Archives of general psychiatry. 2006;63(12):1417–28.

20 Maier S, Tebartz van Elst L, Beier D, Ebert D, Fangmeier T, Radtke M, et al. Increased hippocampal volumes in adults with high functioning autism spectrum disorder and an IQ>100: A manual morphometric study. Psychiatry research. 2015;234(1):152–5.

21 Nicolson R, DeVito TJ, Vidal CN, Sui Y, Hayashi KM, Drost DJ, et al. Detection and mapping of hippocampal abnormalities in autism. Psychiatry research. 2006;148(1):11–21.

22 Haznedar MM, Buchsbaum MS, Hazlett EA, LiCalzi EM, Cartwright C, Hollander E. Volumetric analysis and three-dimensional glucose metabolic mapping of the striatum and thalamus in patients with autism spectrum disorders. The American journal of psychiatry. 2006;163(7):1252–63.

23 Hollander E, Anagnostou E, Chaplin W, Esposito K, Haznedar MM, Licalzi E, et al. Striatal volume on magnetic resonance imaging and repetitive behaviors in autism. Biological psychiatry. 2005;58(3):226–32.

24 Schuetze M, Park MTM, Cho IYK, MacMaster FP, Chakravarty MM, Bray SL. Morphological Alterations in the Thalamus, Striatum, and Pallidum in Autism Spectrum Disorder. Neuropsychopharmacology. 2016;41:2627.

25 Lange N, Travers BG, Bigler ED, Prigge MBD, Froehlich AL, Nielsen JA, et al. Longitudinal volumetric brain changes in autism spectrum disorder ages 6-35 years. Autism Res. 2015;8(1):82–93.

26 McAlonan GM, Suckling J, Wong N, Cheung V, Lienenkaemper N, Cheung C, et al. Distinct patterns of grey matter abnormality in high-functioning autism and Asperger’s syndrome. Journal of child psychology and psychiatry, and allied disciplines. 2008;49(12):1287–95.

27 Lin HY, Ni HC, Lai MC, Tseng WI, Gau SS. Regional brain volume differences between males with and without autism spectrum disorder are highly age-dependent. Molecular autism. 2015;6:29.

28 Tsatsanis KD, Rourke BP, Klin A, Volkmar FR, Cicchetti D, Schultz RT. Reduced thalamic volume in high-functioning individuals with autism. Biological psychiatry. 2003;53(2):121–9.

29 van Rooij D, Anagnostou E, Arango C, Auzias G, Behrmann M, Busatto GF, et al. Cortical and Subcortical Brain Morphometry Differences Between Patients With Autism Spectrum Disorder and Healthy Individuals Across the Lifespan: Results From the ENIGMA ASD Working Group. The American journal of psychiatry. 2018;175(4):359–69.

30 Fair DA, Bathula D, Nikolas MA, Nigg JT. Distinct neuropsychological subgroups in typically developing youth inform heterogeneity in children with ADHD. Proceedings of the National Academy of Sciences of the United States of America. 2012;109(17):6769–74.

31 Li T, van Rooij D, Mota NR, Buitelaar JK, Hoogman M, Vasquez AA, et al. Characterizing neuroanatomic heterogeneity in people with and without ADHD based on subcortical brain volumes. bioRxiv. 2019:868414.

32 Newman ME. Modularity and community structure in networks. Proceedings of the National Academy of Sciences of the United States of America. 2006;103(23):8577–82.

33 Rubinov M, Sporns O. Weight-conserving characterization of complex functional brain networks. NeuroImage. 2011;56(4):2068–79.

34 Karrer B, Levina E, Newman ME. Robustness of community structure in networks. Phys Rev E Stat Nonlin Soft Matter Phys. 2008;77(4 Pt 2):046119.

35 Wen W, Thalamuthu A, Mather KA, Zhu W, Jiang J, de Micheaux PL, et al. Distinct Genetic Influences on Cortical and Subcortical Brain Structures. Scientific Reports. 2016;6:32760.

36 Eyler LT, Prom-Wormley E, Fennema-Notestine C, Panizzon MS, Neale MC, Jernigan TL, et al. Genetic patterns of correlation among subcortical volumes in humans: results from a magnetic resonance imaging twin study. Hum Brain Mapp. 2011;32(4):641–53.

37 Salgado S, Kaplitt MG. The Nucleus Accumbens: A Comprehensive Review. Stereotactic and functional neurosurgery. 2015;93(2):75–93.

38 Sussman D, Leung RC, Chakravarty MM, Lerch JP, Taylor MJ. The developing human brain: age-related changes in cortical, subcortical, and cerebellar anatomy. Brain Behav. 2016;6(4):e00457–e57.

39 Marquand AF, Rezek I, Buitelaar J, Beckmann CF. Understanding Heterogeneity in Clinical Cohorts Using Normative Models: Beyond Case-Control Studies. Biological psychiatry. 2016;80(7):552–61.

40 Coupe P, Catheline G, Lanuza E, Manjon JV. Towards a unified analysis of brain maturation and aging across the entire lifespan: A MRI analysis. Hum Brain Mapp. 2017;38(11):5501–18.

41 Lombardo MV, Lai M-C, Baron-Cohen S. Big data approaches to decomposing heterogeneity across the autism spectrum. Molecular psychiatry. 2019;24(10):1435–50.

42 Ritchie SJ, Cox SR, Shen X, Lombardo MV, Reus LM, Alloza C, et al. Sex Differences in the Adult Human Brain: Evidence from 5216 UK Biobank Participants. Cereb Cortex. 2018;28(8):2959–75.

43 Ecker C, Andrews DS, Gudbrandsen CM, Marquand AF, Ginestet CE, Daly EM, et al. Association Between the Probability of Autism Spectrum Disorder and Normative Sex-Related Phenotypic Diversity in Brain Structure. JAMA Psychiatry. 2017;74(4):329–38.

44 Lai MC, Lerch JP, Floris DL, Ruigrok AN, Pohl A, Lombardo MV, et al. Imaging sex/gender and autism in the brain: Etiological implications. J Neurosci Res. 2017;95(1-2):380–97.

