## Supplementary for "Dissecting the heterogeneous subcortical brain volume of Autism spectrum disorder (ASD) using community detection"

**Table S1. Additional information of methods in each participating site.**

| Cohorts | N (boys/adol/adult) | Age (SD) | Scanner Type | Field strength |
|---|---|---|---|---|
| **ABIDE_CALTECH** | NA / 8 / 18 | NA / 20.3 (1.3) / 34.7 (11.3) | Siemens Trio | 3T |
| **ABIDE_KKI** | 18 / NA / NA | 10.4 (1.3) / NA / NA | Philips Achieva | 3T |
| **ABIDE_LEUVEN_1** | NA / 17 / 10 | NA / 20.4 (1.6) / 26.3 (3.2) | Philips Achieva | 3T |
| **ABIDE_LEUVEN_2** | 18 / 8 / NA | 13.5 (1.0) / 16.1 (0.7) / NA | Philips Achieva | 3T |
| **ABIDE_MAX_MUN** | 13 / 4 / 27 | 10.1 (1.8) / 20.8 (1.6) / 33.0 (9.4) | Siemens Verio | 3T |
| **ABIDE_NYU** | 87 / 35 / 25 | 10.9 (2.2) / 17.7 (1.9) / 26.4 (4.2) | Siemens Allegra | 3T |
| **ABIDE_OHSU** | 18 / NA / NA | 10.3 (1.2) / NA / NA | Siemens Trio | 3T |
| **ABIDE_OLIN** | 8 / 20 / 1 | 12.8 (1.2) / 18.1 (2.2) / 23.0 (0.0) | Siemens Allegra | 3T |
| **ABIDE_PITT** | 16 / 17 /13 | 12.7 (1.6) / 18.4 (2.1) / 29.3 (4.5) | Siemens Allegra | 1.5T |
| **ABIDE_SBL** | NA / 2 / 27 | NA / 21.0 (1.4) / 35.6 (8.1) | Philips Achieva | 3T |
| **ABIDE_ SDSU** | 16 / 13 / 1 | 13.5 (1.0) / 16.2 (0.6) / 37.7 (0.0) | GE MR750 | 3T |
| **ABIDE_STANFORD** | 32 / NA / NA | 10.1 (1.6) / NA / NA | GR Signa | 3T |
| **ABIDE_TCD** | 22 / 28 / 4 | 12.9 (1.5) / 18.3 (2.1) / 24.9 (1.3) | Philips Achieva | 3T |
| **ABIDE_UM_1** | 62 / 23 / NA | 11.3 (1.7) / 16.8 (1.5) / NA | GR Signa | 3T |
| **ABIDE_UM_2** | 16 / 12 / 1 | 13.7 (1.1) / 16.5 (0.9) / 26.8 (0.0) | GR Signa | 3T |
| **ABIDE_USM** | 18 / 38 / 39 | 11.6 (2.3) / 17.9 (1.7) / 29.3 (6.2) | Siemens Trio | 3T |
| **ABIDE_YALE** | 29 / 8 / NA | 11.3 (2.2) / 16.7 (0.9) / N | Siemens Magnetom | 3T |
| **ABIDEII-BNI_1** | NA / 15 / 36 | NA / 19.9 (1.6) / 48.6 (9.7) | Philips Ingenia | 3T |
| **ABIDEII-EMC_1** | 41/ NA / NA | 8.2 (1.1) / NA / NA | GE MR750 | 3T |
| **ABIDEII-ETH_1** | 1 / 13 / 19 | 14.8 (0) / 19.2 (1.7) / 26.1 (2.7) | Philips Ingenia | 3T |
| **ABIDEII-GU_1** | 68 / NA / NA | 10.9 (1.6) / NA / NA | Siemens TriTim | 3T |
| **ABIDEII-IP_1** | 7 / 7 / 9 | 11.1 (2.5) / 17.2 (2.4) / 28.9 (6.8) | Siemens TriTim | 1.5T |
| **ABIDEII-IU_1** | NA / 16 / 12 | NA / 19.9 (1.5) / 28.3 (5.2) | Philips Achieva | 3T |
| **ABIDEII-KKI_1** | 130 / NA / NA | 10.5 (1.4) / NA / NA | Philips Achieva | 3T |
| **ABIDEII-KUL_3** | NA / 11 / 15 | NA / 19.4 (1.5) / 26.9 (4.0) | Philips Achieva | 3T |
| **ABIDEII-NYU_1** | 64 / 1 / 2 | 8.9 (2.2) / 17.9 (0.0) / 26.3 (0.4) | Siemens Allegra | 3T |
| **ABIDEII-NYU_2** | 20 / NA / NA | 6.8 (1.0) / NA / NA | Siemens Allegra | 3T |
| **ABIDEII-OHSU_1** | 54 / 18 / NA | 11.2 (2.0) / 15.0 (0.0) / NA | Siemens Skyra | 3T |
| **ABIDEII-OILH_2** | NA /18 / 13 | NA / 19.5 (1.3) / 26.8 (2.4) | Siemens TriTim | 3T |
| **ABIDEII-SDSU_1** | 33 / 14 / NA | 11.1 (2.0) / 16.8 (1.0) / NA | GE MR750 | 3T |
| **ABIDEII-TCD_1** | 19 / 19 / NA | 12.4 (1.5) / 17.9 (1.7) / NA | Philips Achieva | 3T |
| **ABIDEII-UCD_1** | 11 / 11 / NA | 13.2 (1.0) / 16.6 (0.8) / NA | Siemens TriTim | 3T |
| **ABIDEII-UCLA_1** | 23 / 1 / NA | 11.1 (2.2) / 15.0 (0.0) / NA | Siemens TriTim | 3T |

| | | | | |
|---|---|---|---|---|
| **ABIDEII-USM_1** | 8 / 5 / 10 | 12.9 (2.0) / 17.5 (2.0) / 28.2 (5.1) | Siemens TriTim | 3T |
| **Barcelona** | 50 / 19 / NA | 10.7 (2.3) / 15.9 (0.5) / NA | NA | NA |
| **BRC** | 25 / 27 / NA | 12.5 (1.2) / 16.6 (1.2) / NA | GE Signa HDx | NA |
| **CMU** | NA / 6 / 13 | NA / 21.0 (0.6) / 29.6 (5.4) | Siemens Magnetom | 3T |
| **Dresden** | NA / 2 / 16 | NA / 21.6 (0.6) / 36.0 (12.1) | NA | NA |
| **FAIR** | 54 / 2 / NA | 11.4 (1.8) / 15.2 (0.1) / NA | Siemens Trio | 3T |
| **FRANKFURT** | NA / 21 / NA | NA / 18.0 (0.0) / NA | Siemens | 1.5T |
| **FSM** | 40 / NA / NA | 4.1 (0.9) / NA / NA | GE Signa | 1.5T |
| **MRC** | NA / 38 / 106 | NA / 19.7 (1.5) / 29.6 (6.0) | GE Signa HDx | 3T |
| **MYAD** | 43 / NA / NA | 4.7 (1.7) / NA / NA | Siemens symphony | 1.5T |
| **Nijmegen 1** | 15 / 21 / NA | 13.5 (0.9) / 16.5 (1.1) / NA | Siemens Avanto | 1.5T |
| **Nijmegen 2** | NA / 13 / 28 | NA / 20.0 (1.4) / 30.6 (5.1) | NA | NA |
| **Nijmegen 3** | 81 / NA / NA | 9.5 (1.8) / NA / NA | NA | NA |
| **HGGM** | 52 / 11 / NA | 11.6 (2.0) / 16.5 (1.0) / NA | Philips Intera | 1.5T |
| **PITT** | 53 / 48 / 21 | 12.1 (1.8) / 17.7 (2.2) / 28.1 (4.9) | Siemens Allegra | 3T |
| **SAOPAULO** | 13 / 7 / NA | 11.0 (2.5) / 17.3 (1.4) / NA | Philips | 3T |
| **TCD** | 42 / 45 / 1 | 12.9 (1.4) / 18.1 (1.8) / 24.8 (0.0) | Philips Achieva | 3T |
| **TORONTO** | 142 / 39 / NA | 10.4 (2.8) / 16.5 (1.1) / NA | Siemens Trio | 3T |
| **UMCU** | 43 / 16 / 8 | 11.0 (1.9) / 18.8 (1.7) / 23.2 (1.2) | Philips | 1.5T |
| **Total** | 1,505 / 681 / 477 | 10.6 (2.7) / 18.0 (2.0) / 31.2 (8.7) | NA | NA |

**Table S2. The distribution of boys (ASD patients/controls) in each community**

| Cohorts | Patients- Community (%) | | | | | Controls-Community (%) | | | |
|---|---|---|---|---|---|---|---|---|---|
| | 1 | 2 | 3 | 4 | | 1 | 2 | 3 | 4 |
| **ABIDE_KKI** | 2 (28.6%) | 1 (14.3%) | 2 (28.6%) | 2 (28.6%) | | 3 (27.3%) | 2 (18.2%) | 3 (27.3%) | 3 (27.3%) |
| **ABIDE_LEUVEN_2** | 6 (54.5%) | 0 | 4 (36.4%) | 1 (9.1%) | | 2 (28.6%) | 1 (14.3%) | 3 (42.9%) | 1 (14.3%) |
| **ABIDE_MAX_MUN** | 0 | 2 (25.0%) | 5 (62.5%) | 1 (12.5%) | | 0 | 2 (40.0%) | 2 (40.0%) | 1 (20.0%) |
| **ABIDE_NYU** | 6 (13.0%) | 8 (17.4%) | 24 (52.2%) | 8 (17.4%) | | 25 (61.0%) | 1 (2.4%) | 8 (19.5%) | 7 (17.1%) |
| **ABIDE_OHSU** | 5 (71.4%) | 0 | 0 | 2 (28.6%) | | 4 (36.4%) | 1 (9.1%) | 5 (45.5%) | 1 (9.1%) |
| **ABIDE_OLIN** | 2 (50.0%) | 1 (25.0%) | 1 (25.0%) | 0 | | 2 (50.0%) | 0 | 0 | 2 (50.0%) |
| **ABIDE_PITT** | 1 (12.5%) | 2 (25.0%) | 5 (62.5%) | 0 | | 5 (62.5%) | 2 (25.0%) | 1 (12.5%) | 0 |
| **ABIDE_SDSU** | 4 (66.7%) | 1 (16.7%) | 0 | 1 (16.7%) | | 1 (10.0%) | 1 (10.0%) | 0 | 8 (80.0%) |
| **ABIDE_STANFORD** | 2 (12.5%) | 2 (12.5%) | 0 | 12 (75.0%) | | 2 (12.5%) | 2 (12.5%) | 3 (18.8%) | 9 (56.3%) |
| **ABIDE_TCD** | 2 (28.6%) | 0 | 0 | 5 (71.4%) | | 2 (13.3%) | 2 (13.3%) | 4 (26.7%) | 7 (46.7%) |
| **ABIDE_UM_1** | 7 (20.0%) | 6 (17.1%) | 6 (17.1%) | 16 (45.7%) | | 11 (40.7%) | 3 (11.1%) | 6 (22.2%) | 7 (46.7%) |
| **ABIDE_UM_2** | 4 (44.4%) | 0 | 1 (11.1%) | 4 (44.4%) | | 3 (42.9%) | 0 | 0 | 4 (57.1%) |
| **ABIDE_USM** | 1 (11.1%) | 3 (33.3%) | 3 (33.3%) | 2 (22.2%) | | 3 (33.3%) | 2 (22.2%) | 1 (11.1%) | 3 (33.3%) |
| **ABIDE_YALE** | 2 (13.3%) | 6 (40.0%) | 7 (46.7%) | 0 | | 0 | 11 (78.6%) | 2 (14.3%) | 1 (7.1%) |
| **ABIDEII-EMC_1** | 10 (47.6%) | 2 (9.5%) | 7 (33.3%) | 2 (9.5%) | | 12 (60.0%) | 0 | 5 (25.0%) | 3 (15.0%) |
| **ABIDEII-ETH_1** | 1 (100.0%) | 0 | 0 | 0 | | NA | NA | NA | NA |
| **ABIDEII-GU_1** | 7 (17.1%) | 18 (43.9%) | 5 (12.2%) | 11 (26.8%) | | 3 (11.1%) | 15 (55.6%) | 7 (25.9%) | 2 (7.4%) |
| **ABIDEII-IP_1** | 0 | 3 (50.0%) | 3 (50.0%) | 0 | | 0 | 1 (100.0%) | 0 | 0 |
| **ABIDEII-KKI_1** | 5 (14.3%) | 7 (20.0%) | 1 (2.9%) | 22 (62.9%) | | 13 (13.7%) | 14 (14.7%) | 1 (1.1%) | 67 (70.5%) |
| **ABIDEII-NYU_1** | 9 (25.0%) | 5 (13.9%) | 0 | 22 (62.9%) | | 6 (21.4%) | 4 (14.3%) | 1 (3.6%) | 17 (60.7%) |
| **ABIDEII-NYU_2** | 8 (40.0%) | 2 (10.0%) | 2 (10.0%) | 8 (40.0%) | | NA | NA | NA | NA |
| **ABIDEII-OHSU_1** | 5 (18.5%) | 4 (14.8%) | 17 (63.0%) | 1 (3.7%) | | 12 (44.4%) | 3 (11.1%) | 9 (33.3%) | 3 (11.1%) |
| **ABIDEII-SDSU_1** | 9 (47.4%) | 3 (15.8%) | 4 (21.1%) | 3 (15.8%) | | 4 (28.6%) | 1 (7.1%) | 7 (50.0%) | 2 (14.3%) |
| **ABIDEII-TCD_1** | 9 (90.0%) | 0 | 0 | 1 (10.0%) | | 6 (66.7%) | 0 | 0 | 3 (33.3%) |
| **ABIDEII-UCD_1** | 0 | 6 (85.7%) | 1 (14.3%) | 0 | | 0 | 3 (75.0%) | 1 (25.0%) | 0 |
| **ABIDEII-UCLA_1** | 4 (30.8%) | 2 (15.4%) | 1 (7.7%) | 6 (46.2%) | | 3 (30.0%) | 0 | 3 (30.0%) | 4 (40.0%) |
| **ABIDEII-USM_1** | 3 (50.0%) | 1 (16.7%) | 1 (16.7%) | 1 (16.7%) | | 1 (50%) | 0 | 0 | 1 (50.0%) |

| | | | | | | | | | |
|---|---|---|---|---|---|---|---|---|---|
| **Barcelona** | 3 (11.1%) | 3 (11.1%) | 21 (77.8%) | 0 (0.0%) | | 5 (21.7%) | 0 | 17 (73.9%) | 1 (4.3%) |
| **BRC** | 6 (75.0%) | 1 (12.5%) | 0 | 1 (12.5%) | | 9 (52.9%) | 1 (5.9%) | 1 (5.9%) | 6 (35.3%) |
| **FAIR** | 4 (14.8%) | 13 (48.1%) | 5 (18.5%) | 5 (18.5%) |  | 4 (14.8%) | 9 (33.3%) | 4 (14.8%) | 10 (37.0%) |
| **FSM** | 4 (20.0%) | 7 (35.0%) | 5 (25.0%) | 4 (20.0%) | | 6 (30.0%) | 2 (10.0%) | 4 (20.0%) | 8 (40.0%) |
| **MYAD** | 1 (3.1%) | 21 (65.6%) | 3 (9.4%) | 7 (21.9%) | | 1 (9.1%) | 7 (63.6%) | 1 (9.1%) | 2 (18.2%) |
| **Nijmegen 1** | 0 | 6 (85.7%) | 0 (0.0%) | 1 (14.3%) | | 0 | 8 (100.0%) | 0 | 0 |
| **Nijmegen 3** | 2 (5.4%) | 16 (43.2%) | 15 (40.5%) | 4 (10.8%) | | 7 (15.9%) | 15 (34.1%) | 20 (45.5%) | 2 (4.5%) |
| **HGGM** | 0 | 20 (71.4%) | 8 (28.6%) | 0 | | 0 (0.0%) | 17 (70.8%) | 7 (29.2%) | 0 |
| **PITT** | 3 (14.3%) | 7 (33.3%) | 7 (33.3%) | 4 (19.0%) | | 5 (15.6%) | 18 (56.3%) | 3 (9.4%) | 6 (18.8%) |
| **SAOPAULO** | 3 (60.0%) | 0 | 2 (40.0%) | 0 | | 5 (62.5%) | 1 (12.5%) | 2 (25.0%) | 0 |
| **TCD** | 15 (65.2%) | 2 (8.7%) | 3 (13.0%) | 3 (13.0%) | | 10 (52.6%) | 0 | 6 (31.6%) | 3 (15.8%) |
| **TORONTO** | 35 (43.2%) | 19 (23.5%) | 17 (21.0%) | 10 (12.3%) | | 15 (24.6%) | 17 (27.9%) | 16 (26.2%) | 13 (21.3%) |
| **UMCU** | 0 | 22 (84.6%) | 4 (15.4%) | 0 | | 1 (5.9%) | 14 (82.4%) | 2 (11.8%) | 0 (0.0%) |
| **Total** | **190 (24.6%)** | **222 (28.8%)** | **190 (24.6%)** | **170 (22.0%)** | | **191 (26.1%)** | **180 (24.6%)** | **155 (21.1%)** | **207 (28.2%)** |

***Note:*** CD in boys (ASD patients and healthy controls). NA: not available.

**Table S3. The distribution of male adolescents (patients/controls) in each community**

| Cohorts | Patients- Community (%) | | | | Controls- Community (%) | | |
|---|---|---|---|---|---|---|---|
| | 1 | 2 | 4 | | 1 | 2 | 4 |
| **ABIDE_CALTECH** | 3 (60.0%) | 1 (20.0%) | 1 (20.0%) | | 1 (33.3%) | 0 | 2 (66.7%) |
| **ABIDE_ LEUVEN_1** | 2 (20.0%) | 4 (40.0%) | 4 (40.0%) | | 4 (57.1%) | 2 (28.6%) | 1 (14.3%) |
| **ABIDE_LEUVEN_2** | 1 (100.0%) | 0 | 0 | | 4 (57.1%) | 2 (28.6%) | 1 (14.3%) |
| **ABIDE_MAX_MUN** | 0 | 1 (50.0%) | 1(50.0%) | | 1 (50.0%) | 1 (50.0%) | 0 |
| **ABIDE_NYU** | 4 (40.0%) | 5 (50.0%) | 1 (10.0%) | | 15 (60.0%) | 7 (28.0%) | 3 (12.0%) |
| **ABIDE_OLIN** | 7 (53.8%) | 1 (7.7%) | 5 (38.4%) | | 3 (42.9%) | 0 | 4 (57.1%) |
| **ABIDE_PITT** | 3 (30.0%) | 7 (70.0%) | 0 | | 5 (71.4%) | 1 (14.3%) | 1 (14.3%) |
| **ABIDE_SBL** | 1 (100.0%) | 0 | 0 | | 1 (100.0%) | 0 | 0 |
| **ABIDE_SDSU** | 4 (57.1%) | 0 | 3 (42.9%) | | 0 | 0 | 6 (100.0%) |
| **ABIDE_TCD** | 0 | 4 (26.7%) | 11 (73.3%) | | 1 (7.7%) | 2 (15.4%) | 10 (76.9%) |
| **ABIDE_UM_1** | 3 (33.3%) | 3 (33.3%) | 3 (33.3%) | | 2 (14.3%) | 6 (42.9%) | 6 (42.9%) |
| **ABIDE_UM_2** | 2 (40.0%) | 1 (20.0%) | 2 (40.0%) | | 1 (14.3%) | 1 (14.3%) | 5 (71.4%) |
| **ABIDE_USM** | 3 (11.1%) | 22 (81.5%) | 2 (7.4%) | | 3 (27.3%) | 3 (27.3%) | 5 (45.5%) |
| **ABIDE_YALE** | 0 | 5 (100.0%) | 0 | | 0 | 3 (100.0%) | 0 |
| **ABIDEII-ENI_1** | 8 (88.9%) | 1 (11.1%) | 0 | | 4 (66.7%) | 2 (33.3%) | 0 |
| **ABIDEII-ETH_1** | 2 (28.6%) | 2 (28.6%) | 3 42.9%) | | 3 (50.0%) | 0 | 3 (50.0%) |
| **ABIDEII-IU_1** | 0 | 9 (100.0%) | 0 | | 0 | 7 (100.0%) | 0 |
| **ABIDEII-KUL_3** | 6 (54.5%) | 2 (18.2%) | 3 (27.2%) | | NA | NA | NA |
| **ABIDEII-NYU_2** | 0 | 0 | 1 (100.0%) | | NA | NA | NA |
| **ABIDEII-OHSU_1** | 0 | 2 (100.0%) | 0 | | NA | NA | NA |
| **ABIDEII-OILH_2** | 0 | 9 (90.0%) | 1 (10.0%) | | 0 | 8 (100.0%) | 0 |
| **ABIDEII-SDSU_1** | 3 (42.9%) | 3 (42.9%) | 1 (14.3%) | | 4 (57.1%) | 0 | 3 (42.9%) |
| **ABIDEII-TCD_1** | 2 (25.0%) | 1 (12.5%) | 5 (62.5%) | | 6 (54.5%) | 0 | 5 (45.5%) |
| **ABIDEII-UCD_1** | 0 | 0 | 1 (100.0%) | | 0 | 6 (100.0%) | 0 |
| **ABIDEII-USM_1** | 1 (33.3%) | 0 | 2 (66.7%) | | 1 (50.0%) | 0 | 1 (50.0%) |
| **Barcelona** | 3 (25.3%) | 4 (40.0%) | 3 (30.0%) | | 2 (22.2%) | 2 (22.2%) | 5 (55.6%) |
| **BRC** | 6 (54.5%) | 4 (36.4%) | 1 (9.1%) | | 10 (62.5%) | 3 (18.8%) | 3 (18.8%) |
| **CMU** | 1 (33.3%) | 2 (66.7%) | 0 | | 1 (33.3%) | 2 (66.7%) | 0 |
| **Dresden** | 0 | 1 (100.0%) | 0 | | 0 | 1 (100.0%) | 0 |
| **FAIR** | 0 | 2 (100.0%) | 0 | | NA | NA | NA |
| **FRANKFURT** | 1 (10.0%) | 9 (90.0%) | 0 | | 0 | 10 (90.9%) | 1 (9.1%) |

| | | | | | | | |
|---|---|---|---|---|---|---|---|
| **MRC** | 0 | 3 (11.1%) | 24 (88.9%) | | 1 (9.1%) | 0 | 10 (90.9%) |
| **Nijmegen 1** | 0 | 7 (100.0%) | 0 | | 0 | 13 92.8%) | 1 (7.2%) |
| **Nijmegen 2** | 0 | 9 (100.0%) | 0 | | 0 | 4 (100.0%) | 0 |
| **HGGM** | 0 | 5 (100.0%) | 0 | | 0 | 6 (100.0%) | 0 |
| **PITT** | 6 (31.6%) | 7 (36.8%) | 6 (31.6%) | | 3 (20.0%) | 10 (66.7%) | 2 (13.3%) |
| **SAOPAULO** | 2 (50.0%) | 1 (25.5%) | 1 (25.5%) | | 1 (33.3%) | 1 (33.3%) | 1 (33.3%) |
| **TCD** | 8 (38.1%) | 6 (28.6%) | 7 (33.3%) | | 8 (33.3%) | 9 (37.5%) | 7 (29.2%) |
| **TORONTO** | 9 (37.5%) | 11 (45.8%) | 4 (16.7%) | | 4 (28.6%) | 8 (57.1%) | 2 (14.3%) |
| **UMCU** | 0 | 7 (100.0%) | 0 | | 0 | 9 (100.0%) | 0 |
| **Total** | **91 (25.3%)** | **173 (48.1%)** | **96 (26.7%)** | | **93 (29.0%)** | **132 (41.1%)** | **96 (29.9%)** |

***Note****:* CD in male adolescents (ASD patients and healthy controls). NA: not available.

**Table S4. The distribution of adult men (patients/controls) in each community**

| Cohorts | Patients- Community (%) | | | | Controls- Community (%) | | |
|---|---|---|---|---|---|---|---|
| | 1 | 2 | 4 | | 1 | 2 | 4 |
| **ABIDE_CALTECH** | 4 (50.0%) | 4 (50.0%) | 0 | | 1 (10.0%) | 6 (60.0%) | 3 (30.0%) |
| **ABIDE_ LEUVEN_1** | 0 | 3 (75.0%) | 1 (25.0%) | | 2 (33.3%) | 4 (66.7%) | 0 |
| **ABIDE_MAX_MUN** | 3 (30.0%) | 3 (30.0%) | 4 (40.0%) | | 3 (17.6%) | 10 (58.8%) | 4 (23.5%) |
| **ABIDE_NYU** | 6 (60.0%) | 4 (40.0%) | 0 | | 8 (53.3%) | 5 (33.3%) | 2 (13.3%) |
| **ABIDE_OLIN** | NA | NA | NA | | 1 (100.0%) | 0 | 0 |
| **ABIDE_PITT** | 2 (25.0%) | 5 (62.5%) | 1 (12.5%) | | 3 (60.0%) | 2 (40.0%) | 0 |
| **ABIDE_SBL** | 7 (50.0%) | 4 (28.6%) | 3 (21.4%) | | 1 (7.7%) | 10 (76.9%) | 2 (15.4%) |
| **ABIDE_SDSU** | 0 | 0 | 1 (100.0%) | | NA | NA | NA |
| **ABIDE_TCD** | 1 (50.0%) | 1 (50.0%) | 0 | | 1 (50.0%) | 1 (50.0%) | 0 |
| **ABIDE_UM_2** | NA | NA | NA | | 1 (100.0%) | 0 | 0 |
| **ABIDE_USM** | 6 (28.6%) | 9 (42.9%) | 6 (28.6%) | | 6 (33.3%) | 7 (38.9%) | 5 (27.8%) |
| **ABIDEII-ENI_1** | 7 (41.2%) | 8 (47.1%) | 2 (11.8%) | | 4 (21.1%) | 13 (68.4%) | 2 (10.5%) |
| **ABIDEII-ETH_1** | 3 (60.0%) | 1 (20.0%) | 1 (20.0%) | | 5 (35.7%) | 7 (50.0%) | 2 (14.3%) |
| **ABIDEII-IP_1** | 1 (100.0%) | 0 | 0 | | 3 (37.5%) | 3 (37.5%) | 2 (25.0%) |
| **ABIDEII-IU_1** | 2 (40.0%) | 3 (60.0%) | 0 | | 4 (57.1%) | 3 (42.9%) | 0 |
| **ABIDEII-KUL_3** | 8 (53.3%) | 6 (40.0%) | 1 (6.7%) | | NA | NA | NA |
| **ABIDEII-NYU_2** | 0 | 2 (100.0%) | 0 | | NA | NA | NA |
| **ABIDEII-OILH_2** | 3 (75.0%) | 1 (25.0%) | 0 | | 3 (33.3%) | 5 (55.6%) | 1 (11.1%) |
| **ABIDEII-USM_1** | 1 (50.0%) | 0 | 1 (50.0%) | | 4 (50.0%) | 4 (50.0%) | 0 |
| **CMU** | 2 (33.3%) | 2 (33.3%) | 2 (33.3%) | | 0 | 5 (71.4%) | 2 (2.6%) |
| **Dresden** | 5 (35.7%) | 8 (57.1%) | 1 (7.1%) | | 0 | 2 | 0 |
| **MRC** | 10 (22.7%) | 15 (34.1%) | 26 (43.3%) | | 12 (19.4%) | 27 (43.5%) | 23 (37.1%) |
| **Nijmegen 3** | 2 (11.8%) | 13 (76.5%) | 2 (11.8%) | | 2 (18.2%) | 8 (72.7%) | 1 (9.1%) |
| **PITT** | 0 | 5 (62.5%) | 3 (37.5%) | | 2 (28.6%) | 3 (42.9%) | 2 (28.6%) |
| **UMCU** | 0 | 2 (100.0%) | 0 | | 0 | 5 (83.1%) | 1 (16.7%) |
| **TCD** | NA | NA | NA | | 0 | 1 (100.0%) | 0 |
| **Total** | **73 (33.0%)** | **100 (45.2%)** | **48 (21.7%)** | | **69 (27.2%)** | **132 (52.0%)** | **53 (20.9%)** |

***Note:*** CD in adult men (ASD patients and healthy controls). NA: not available.

**Table S5. Comparison of subcortical brain volumes within each community and the whole subsample of boys**

| | **Total sample** | | | | **Community 1** | | | | **Community 2** | | | |
|---|---|---|---|---|---|---|---|---|---|---|---|---|
| | **Mean volume Patients/Controls** | ***p value*** | ***adjusted p value*** | **Cohen's d (95% CIs)** | **Mean volume Patients/Controls** | ***p value*** | ***adjusted p value*** | **Cohen's d (95% CIs)** | **Mean volume Patients/Controls** | ***p value*** | ***adjusted p value*** | **Cohen's d (95% CIs)** |
| **Accumbens** | 697.1 (140.2)/ 712.1 (133.9) | 0.08 | 0.24 | -0.10 (-0.20 - 2.0e-03) | 804.8 (113.8)/ 812.0 (100.0) | 0.80 | 0.90 | -0.06 (-0.25 - 0.13) | 594.5 (101.2)/ 600.2 (109.6) | 0.57 | 0.74 | -0.02 (-0.20 - 0.17) |
| **Caudate** | 4125.3 (561.2)/ 4138.9 (553.7) | 0.66 | 0.81 | -0.02 (-0.12 - 0.08) | 4206.3 (524.2)/ 4250.8 (523.1) | 0.66 | 0.81 | -0.06 (-0.26 - 0.13) | 4050.0 (580.9)/ 4015.7 (547.5) | 0.22 | 0.45 | 0.14 (-0.05 - 0.32) |
| **Putamen** | 6332.8 (795.2)/ 6387.3 (817.6) | 0.31 | 0.51 | -0.05 (-0.15 - 0.05) | 6754.9 (666.5)/ 6851.6 (585.8) | 0.14 | 0.35 | -0.13 (-0.33 - 0.06) | 5922.9 (667.2)/ 5843.6 (703.2) | 0.11 | 0.30 | 0.20 (0.01 - 0.39) |
| **Pallidum** | 1855.6 (270.5)/ 1866.7 (274.5) | 0.41 | 0.62 | -0.04 (-0.15 - 0.06) | 1892.0 (231.5)/ 1890.0 (225.2) | 0.86 | 0.93 | 0.02 (-0.17 -0.21) | 1860.5 (276.0)/ 1802.1 (278.3) | **5.0e-03** | **0.03** | 0.29 (0.10 – 0.48) |
| **Amygdala** | 1638.8 (270.7)/ 1659.09 (256.3) | 0.14 | 0.35 | -0.06 (-0.17 - 0.04) | 1803.5 (251.2)/ 1780.2 (248.1) | 0.35 | 0.55 | 0.17 (-0.03 - 0.36) | 1462.9 (246.2)/ 1548.4 (205.4) | **2.0e-05** | **2.0e-04** | -0.37 (-0.56 - -0.19) |
| **Hippocampus** | 4198.0 (564.5)/ 4203.5 (518.5) | 0.84 | 0.92 | 0.01 (-0.09 - 0.11) | 4370.8 (483.1)/ 4263.5 (465.2) | **9.0e-03** | **0.05** | 0.37 (0.18 - 0.57) | 3996.0 (550.3)/ 4238.5 (482.0) | **5.0e-09** | **1.2e-07** | -0.55 (-0.74 - -0.36) |
| **Thalamus** | 7773.0 (879.9)/ 7783.9 (893.9) | 0.68 | 0.81 | 5e-04 (-0.10 - 0.10) | 7493.0 (780.4)/ 7540.3 (779.0) | 0.28 | 0.48 | -0.02 (-0.22 - 0.17) | 8069.5 (983.6)/ 8204.1 (891.5) | 0.28 | 0.48 | -0.09 (-0.27 – 0.10) |

Continued

| | Community 3 | | | | Community 4 | | | |
|---|---|---|---|---|---|---|---|---|
| | **Mean volume Patients/Controls** | ***p value*** | ***adjusted p value*** | **Cohen's d (95% CIs)** | **Mean volume Patients/Controls** | ***p value*** | ***adjusted p value*** | **Cohen's d (95% CIs)** |
| **Accumbens** | 633.0 (110.4)/ 714.8 (100.3) | **2.8e-11** | **8.8e-10** | -0.84 (-1.07 - -0.60) | 758.8 (108.5)/ 751.8 102.5) | 0.25 | 0.46 | 0.13 (-0.09 - 0.35) |
| **Caudate** | 3922.1 (511.3)/ 3937.1 (531.1) | 0.10 | 0.29 | -0.21 (-0.44 - 0.01) | 4363.7 (524.1)/ 4341.3 (520.3) | 0.38 | 0.59 | 0.11 (-0.11 - 0.33) |
| **Putamen** | 5869.4 (615.3)/ 6078.1 (668.3) | **2.3e-05** | **2.0e-04** | -0.50 (-0.73 - -0.27) | 6850.7 (678.3)/ 6859.7 (712.5) | 0.77 | 0.89 | 0.09 (-0.13 - 0.31) |
| **Pallidum** | 1696.2 (241.3)/ 1721.0 (237.2) | 0.05 | 0.17 | -0.23 (-0.46 - -0.01) | 1996.8 (252.1)/ 2047.5 (249.3) | 0.08 | 0.24 | -0.19 (-0.42 - 0.03) |
| **Amygdala** | 1691.8 (204.1)/ 1768.6 (270.3) | **9.0e-07** | **1.2e-05** | -0.61 (-0.84 - -0.38) | 1568.8 (238.7)/ 1587.3 (223.5) | 0.94 | 0.99 | 0.05 (-0.18 - 0.26) |
| **Hippocampus** | 4481.1 (497.2)/ 4479.2 (504.6) | **0.02** | 0.10 | -0.28 (-0.51 - -0.06) | 3862.1 (496.6)/ 3862.1 (467.7) | 0.25 | 0.46 | 0.15 (-0.07 - 0.38) |
| **Thalamus** | 7916.0 (815.5)/ 7518.1 (800.4) | **4.2e-05** | **3.2e-04** | 0.37 (0.14 - 0.59) | 7602.7 (788.6)/ 7675.0 (880.0) | 0.97 | 1.00 | 0.01 (-0.21 - 0.23) |

***Note:*** Adjusted p value: FDR correction in subcortical volumes across age bins. Significant difference in bold. 95% CIs: 95% Confidence intervals.

**Table S6. Comparison of subcortical brain volumes within each community and the whole subsample of male adolescents**

| | Total sample | | | | Community 1 | | | |
|---|---|---|---|---|---|---|---|---|
| | **Mean volume Patients/Controls** | ***p value*** | ***adjusted p value*** | **Cohen's d (95% CIs)** | **Mean volume Patients/Controls** | ***p value*** | ***adjusted p value*** | **Cohen's d (95% CIs)** |
| **Accumbens** | 681.4 (142.9)/ 716.9 (150.6) | **2.0e-03** | **0.01** | -0.24 (-0.39 - -0.09) | 756.3 (116.9)/ 818.6 (126.6) | **1.4e-05** | **1.6e-04** | -0.54 (-0.82 - -0.27) |
| **Caudate** | 4103.4 (562.2)/ 4184.3 (548.1) | 0.05 | 0.17 | -0.15 (-0.30 - -3.0e-03) | 4093.6 (507.6)/ 4235.7 (521.3) | 0.04 | 0.15 | -0.28 (-0.55 - -0.01) |
| **Putamen** | 6322.5 (802.4)/ 6547.8 (755.8) | **2.3e-05** | **2.0e-04** | -0.29 (-0.44 - -0.13) | 6537.1 (600.3)/ 6964.3 (569.5) | **4.0e-07** | **5.8e-06** | -0.73 (-1.01 - -0.45) |
| **Pallidum** | 1825.6 (288.2)/ 1858.6 (259.6) | 0.12 | 0.31 | -0.12 (-0.27 - 0.03) | 1773.6 (220.8)/ 1869.3 (224.9) | **3.0e-03** | **0.02** | -0.42 (-0.69 - -0.14) |
| **Amygdala** | 1687.2 (273.1)/ 1730.7 (251.1) | **0.03** | 0.12 | -0.17 (-0.32 - -0.02) | 1926.8 (210.3)/ 1922.6 (189.6) | 0.72 | 0.85 | 0.04 (-0.23 - 0.31) |
| **Hippocampus** | 4368.3 (520.9)/ 4442.1 (478.2) | **0.03** | 0.12 | -0.17 (-0.32 - -0.01) | 4639.0 (459.0)/ 4600.4 (458.0) | 0.28 | 0.48 | 0.14 (-0.13 - 0.42) |
| **Thalamus** | 8030.2 (1002.8)/ 8088.4 (897.4) | 0.25 | 0.46 | -0.09 (-0.24 - 0.06) | 7659.0 (804.9)/ 7803.8 (688.8) | 0.20 | 0.42 | -0.16 (-0.44 - 0.11) |

Continued

| | Community 2 | | | | Community 4 | | | |
|---|---|---|---|---|---|---|---|---|
| | **Mean volume Patients/Controls** | ***p value*** | ***adjusted p value*** | **Cohen's d (95% CIs)** | **Mean volume Patients/Controls** | ***p value*** | ***adjusted p value*** | **Cohen's d (95% CIs)** |
| **Accumbens** | 602.4 (124.7)/ 616.7 (118.5) | 0.05 | 0.17 | -0.12 (-0.35 - 0.11) | 730.3 (131.5)/ 766.8 (111.2) | 0.15 | 0.37 | -0.25 (-0.56 - 0.06) |
| **Caudate** | 4030.5 (575.2)/ 4051.6 (539.5) | 0.57 | 0.74 | -0.09 (-0.32 - 0.13) | 4234.8 (579.5)/ 4370.2 (543.0) | 0.34 | 0.54 | -0.19 (-0.50 - 0.12) |
| **Putamen** | 5885.0 (701.5)/ 5981.1 (538.2) | 0.50 | 0.70 | -0.17 (-0.39 - 0.06) | 6812.4 (778.7)/ 7058.5 (586.3) | **0.04** | 0.14 | -0.33 (-0.64 - -0.03) |
| **Pallidum** | 1780.2 (309.1)/ 1768.3 (248.4) | 0.16 | 0.37 | 0.01 (-0.22 - 0.24) | 1958.7 (278.0)/ 2019.9 (249.0) | 0.27 | 0.48 | -0.19 (-0.50 - 0.12) |
| **Amygdala** | 1625.1 (204.8)/ 1654.1 (229.9) | 0.87 | 0.93 | -0.19 (-0.41 - 0.04) | 1528.0 (261.0)/ 1604.7 (206.5) | 0.18 | 0.40 | -0.27 (-0.57 - 0.04) |
| **Hippocampus** | 4395.7 (483.1)/ 4459.7 (468.4) | 0.20 | 0.42 | -0.24 (-0.46 - -0.01) | 4026.3 (453.9)/ 4179.9 (417.1) | 0.11 | 0.30 | -0.30 (-0.60 - 0.01) |
| **Thalamus** | 8560.1 (942.0)/ 8422.9 (938.2) | 0.64 | 0.81 | 0.04 (-0.19 - 0.26) | 7558.6 (876.3)/ 7842.8 (869.8) | 0.180 | 0.40 | -0.28 (-0.59 - 0.03) |

***Note:*** Adjusted p value: FDR correction in subcortical volumes across age bins. Significant difference in bold. 95% CIs: 95% Confidence intervals.

**Table S7. Comparison of subcortical brain volumes within each community and the whole subsample of adult men**

| | Total sample | | | | Community 1 | | | |
|---|---|---|---|---|---|---|---|---|
| | **Mean volume Patients/Controls** | ***p value*** | ***adjusted p value*** | **Cohen's d (95% CIs)** | **Mean volume Patients/Controls** | ***p value*** | ***adjusted p value*** | **Cohen's d (95% CIs)** |
| **Accumbens** | 649.4 (137.9)/ 663.8 (151.9) | 0.23 | 0.45 | -0.11 (-0.29 - 0.07) | 787.4 (132.1)/ 814.3 (126.7) | 0.24 | 0.46 | -0.21 (-0.56 - 0.13) |
| **Caudate** | 4033.8 (538.4)/ 4002.8 (580.5) | 0.67 | 0.81 | 0.04 (-0.14 - 0.22) | 4024.2 (483.9)/ 4080.3 (541.5) | 0.48 | 0.69 | -0.12 (-0.46 - 0.22) |
| **Putamen** | 6228.7 (878.6)/ 6339.8 (812.2) | 0.19 | 0.41 | -0.13 (-0.31 - 0.06) | 6798.2 (801.1)/ 6729.5 (678.3) | 0.48 | 0.69 | 0.13 (-0.21 - 0.47) |
| **Pallidum** | 1726.1 (275.5)/ 1761.8 (265.1) | 0.16 | 0.37 | -0.13 (-0.31 - 0.05) | 1800.8 (262.3)/ 1814.7 (242.8) | 0.78 | 0.89 | -0.05 (-0.39 - 0.30) |
| **Amygdala** | 1668.0 (280.7)/ 1656.5 (295.8) | 0.81 | 0.90 | -0.02 (-0.21 - 0.16) | 1862.3 (245.5)/ 1825.1 (255.7) | 0.47 | 0.69 | 0.11 (-0.23 - 0.45) |
| **Hippocampus** | 4387.9 (578.4)/ 4328.6 (574.0) | 0.73 | 0.85 | 0.03 (-0.15 - 0.21) | 4375.6 (505.0)/ 4412.8 (524.6) | 0.49 | 0.69 | -0.14 (-0.48 - 0.20) |
| **Thalamus** | 7933.8 (1051.7)/ 7897.1 (962.3) | 0.62 | 0.80 | -0.04 (-0.22 - 0.14) | 7516.3 (954.1)/ 7650.3 (830.9) | 0.22 | 0.45 | -0.20 (-0.54 - 0.14) |

Continued

| | Community 2 | | | | Community 4 | | | |
|---|---|---|---|---|---|---|---|---|
| | **Mean volume Patients/Controls** | ***p value*** | ***adjusted p value*** | **Cohen's d (95% CIs)** | **Mean volume Patients/Controls** | ***p value*** | ***adjusted p value*** | **Cohen's d (95% CIs)** |
| **Accumbens** | 586.5 (101.9)/ 576.3 (112.2) | 0.77 | 0.89 | 0.05 (-0.21 - 0.31) | 618.7 (92.2)/ 649.0 (96.0) | 0.11 | 0.30 | -0.31 (-0.68 - 0.06) |
| **Caudate** | 3970.7 (546.6)/ 3840.9 (575.9) | 0.28 | 0.48 | 0.15 (-0.12 - 0.41) | 4159.0 (564.9)/ 4223.8 (551.1) | 0.87 | 0.93 | -3.0e-03 (-0.37 - 0.36) |
| **Putamen** | 5680.5 (667.2)/ 5819.4 (642.4) | 0.03 | 0.12 | -0.30 (-0.57 - -0.03) | 6629.5 (684.8)/ 6884.7 (620.5) | 0.16 | 0.37 | -0.25 (-0.62 - 0.12) |
| **Pallidum** | 1595.9 (241.9)/ 1642.0 (249.4) | 0.03 | 0.12 | -0.29 (-0.55 - -0.02) | 1885.1 (234.2)/ 1933.4 (201.0) | 0.57 | 0.74 | -0.04 (-0.41 - 0.32) |
| **Amygdala** | 1651.2 (252.1)/ 1646.3 (276.4) | 0.49 | 0.69 | -0.09 (-0.35 - 0.18) | 1494.2 (239.5)/ 1465.9 (258.8) | 0.80 | 0.90 | 0.07 (-0.30 - 0.44) |
| **Hippocampus** | 4635.7 (501.4)/ 4451.2 (546.0) | 0.05 | 0.17 | 0.27 (0.00 - 0.53) | 3948.7 (522.3)/ 3980.2 (555.6) | 0.26 | 0.47 | -0.14 (-0.51 - 0.23) |
| **Thalamus** | 8295.0 (1031.3)/ 8216.8 (991.3) | 0.33 | 0.53 | -0.11 (-0.37 - 0.16) | 7714.4 (975.8)/ 7571.6 (865.2) | 0.25 | 0.46 | 0.29 (-0.37 - 0.16) |

***Note:*** Adjusted p value: FDR correction in subcortical volumes across age bins. 95% CIs: 95% Confidence intervals.

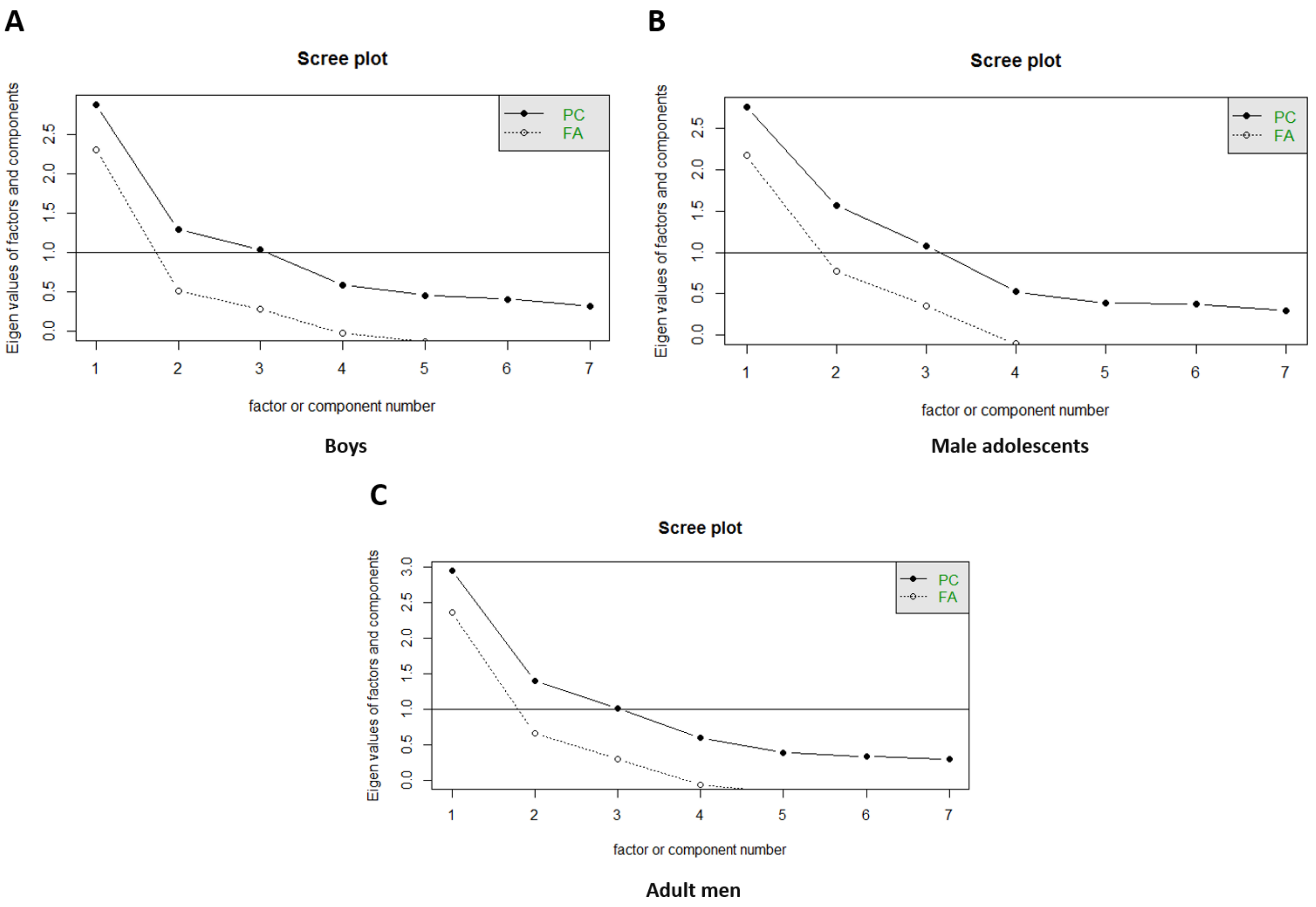


**Figure S1.** Scree plot in each subsample

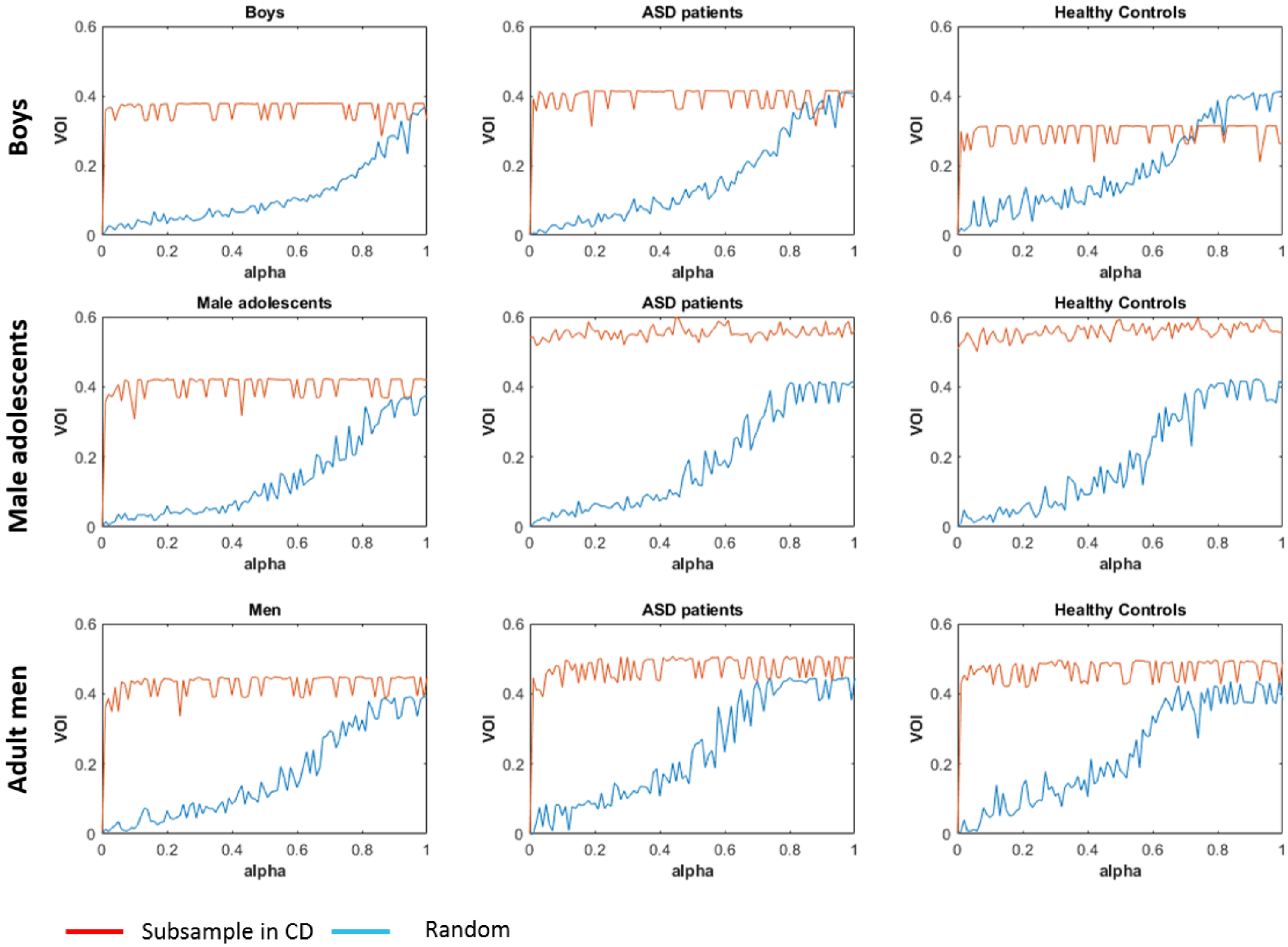


**Figure S2.** VOI in each subsample. VOI: variation of information. alpha: a proportion of edges of a network was randomly perturbed.
